# Genetic and behavioural requirements for structural brain plasticity

**DOI:** 10.1101/431999

**Authors:** Dulcie A Vousden, Alexander Friesen, Xianglan Wen, Lily R Qiu, Nicholas O’Toole, Benjamin C Darwin, Leigh Spencer Noakes, Rylan Allemang Grand, Josie Diorio, Paul W Frankland, Sheena A Josselyn, Brian J Nieman, Michael Meaney, Tie-Yuan Zhang, Jason P Lerch

## Abstract

Human MRI studies show that experience can lead to changes in the volume of task-specific brain regions; however, the behavioural and molecular processes driving these changes remain poorly understood. Here, we used in-vivo mouse MRI and RNA sequencing to investigate the neuroanatomical and transcriptional changes induced by environmental enrichment, exercise, and social interaction. Additionally, we asked whether the volume changes require CREB, a transcription factor critical for memory formation and neuronal plasticity. Enrichment rapidly increased cortical and hippocampal volume, and these effects were not attributable to exercise or social interaction. Instead, they likely arise from learning and sensorimotor experience. Nevertheless, the volume changes were not attenuated in mice with memory impairments caused by loss of CREB, indicating that these effects are driven by processes distinct from this canonical learning and memory pathway. Finally, within brain regions that underwent volume changes, enrichment increased the expression of genes associated with axonogenesis, dendritic spine development, synapse structural plasticity, and neurogenesis, suggesting these processes underlie the volume changes detected with MRI.

---

There is cumulative evidence that experience can leave a lasting impact on brain structure and function [1, 2]. Human neuroimaging studies show that a variety of life experiences can lead to morphological changes to brain anatomy at a scale detectable with MRI. For instance, experiences such as pregnancy [3], early life adversity [4], and psychosocial trauma [5, 6], as well as skill learning, such as learning to juggle [7–9] or how to navigate a complex environment [10–12] have been associated with changes in grey matter volume or density. These studies demonstrate the sensitivity of whole-brain, anatomical MRI to longitudintally monitor experience-dependent plasticity brain in both health and disease. However, the cellular and molecular mechanisms underpinning these mesoscopic changes in anatomy are poorly understood. Indeed, changes in volume have been hypothesized to reflect neuronal remodelling [13], shifts in dendritic spine density [14], neurogenesis, changes in glial number or morphology, angiogenesis, or changes to the extracellular matrix (for review see [15] and [16]). This gap in knowledge considerably limits our ability to interpret the findings observed in human neuroimaging studies.

Rodent neuroimaging studies provide a bridge between molecular neurobiology and human neuroimaging. Rodent MRI is a highly efficient approach to investigate the effects of experience on brain anatomy in a manner that is directly comparable to human neuroimaging studies, but with the added ability to probe the cause of these changes in more depth. In rodents, environmental enrichment is a popular and well-studied experimental model of how experience shapes the brain. Environmental enrichment refers to any laboratory housing environment with supplementary stimulation beyond the standard housing environment, and typically includes tunnels, toys, nesting materials, running wheels, and changes in the amount or potentially quality of social interaction [2, 17]. Environmental enrichment has been shown to have a variety of effects on the brain, including increasing cortical thickness, dendritic complexity and spine density; promoting neurogenesis, gliogenesis, and angiogenesis; and improving learning [1, 2, 18]. *In vivo* rodent MRI has also shown that enrichment increases the volume of cortical and subcortical structures, though whether these changes reflect the same cellular plasticity detected with histological techniques has not been established [19]. Indeed, to our knowledge, no studies to date have attempted to manipulate the putative molecular underpinnings of experiencedependent volume changes.

Despite the long history of environmental enrichment, the particular variables driving the effects of enrichment are incompletely understood. Enrichment includes many factors, including social interaction, increased physical activity, sensorimotor stimulation, novelty of the environment, and complex spatial learning. One hypothesis is that the effects of enrichment on the brain are due to increased general arousal associated with exposure to novel stimuli; others propose that the effects are largely due to increased exercise or voluntary motor activity [2]. Finally, many studies suggest that the effects of enrichment are due to learning and memory formation associated with exploration of a novel, complex environment [2]. In general, no single variable has been shown to account for all the effects of enrichment, but it has been difficult to dissociate the effects of enriched housing from exercise [2]. Moreover, those studies that have endeavoured to isolate the effects of one of these factors examine only a few brain areas, whereas it seems likely that different aspects of enrichment affect certain brain areas more than others. To relate any morphological changes induced by environmental enrichment in rodents to experience-dependent plasticity in the human brain, we need to understand which aspects of the enriched experience are driving which forms of plasticity, and in which brain areas.

Here, we investigated the behavioural and molecular factors underpinning experience-dependent neuroanatomical plasticity in the rodent brain, using whole-brain mouse MRI, RNA sequencing, and a mouseline with impaired learning and memory formation caused by disruption of CREB-dependent transcription. We show that environmental enrichment leads to widespread increases in cortical and hippocampal volume in the adult mouse brain as soon as 2 days after the onset of enrichment, which restricts the potential underlying processes to fast events such as spinogenesis. Secondly, we demonstrate that across the brain, the effects of enrichment on brain anatomy cannot be explained by exercise or social interaction effects alone but instead are due to cognitive stimulation (i.e. exploration and learning about a novel environment) and/or interaction between enrichment factors. Consistent with this, we show that enrichment and exercise drive distinct transcriptional programmes. Thirdly, we find that blocking memory formation via loss of the transcription factor CREB does not attenuate the effects of enrichment on brain anatomy. This indicates that the volume changes are driven by pathways distinct from this canonical learning and memory signalling pathway. Finally, we find evidence that environmental enrichment increases the expression of gene sets associated with positive regulation of axonogenesis, dendritic spine development, synapse structural plasticity, and neurogenesis, and leads to differential expression of genes that regulate angiogenesis; conversely, genes associated with regulation of gliogenesis and the extracellular matrix are not differentially expressed following enrichment. This data suggests axonogenesis, dendritic spine plasticity, and neurogenesis underlie the neuroanatomical changes associated with enrichment.

## 2 Results and Discussion

### 2.1 Environmental enrichment has widespread effects on adult neuroanatomy

We first characterized the effects of a short period of environmental enrichment on mouse neuroanatomy. To do so, we used *in vivo*, high-field MRI to longitudinally image the brains of wildtype (B6/129) mice at baseline and then after 2, 8, and 16 days of housing in an enriched or standard housing environment [20, 21]. We then used automated image registration and segmentation tools to quantify neuroanatomical volume changes for each subject. Consistent with prior studies[19], environmental enrichment had widespread effects on neuroanatomy and was associated with a significant (FDR (false discovery rate) < 10%) change in the growth rate of 55 structures and voxels throughout the cortex and hippocampus, relative to standard-housed control animals (Figures 1, 2, S2, S3, see Supplementary Tables 6-8 for full data).

**Figure 1:**
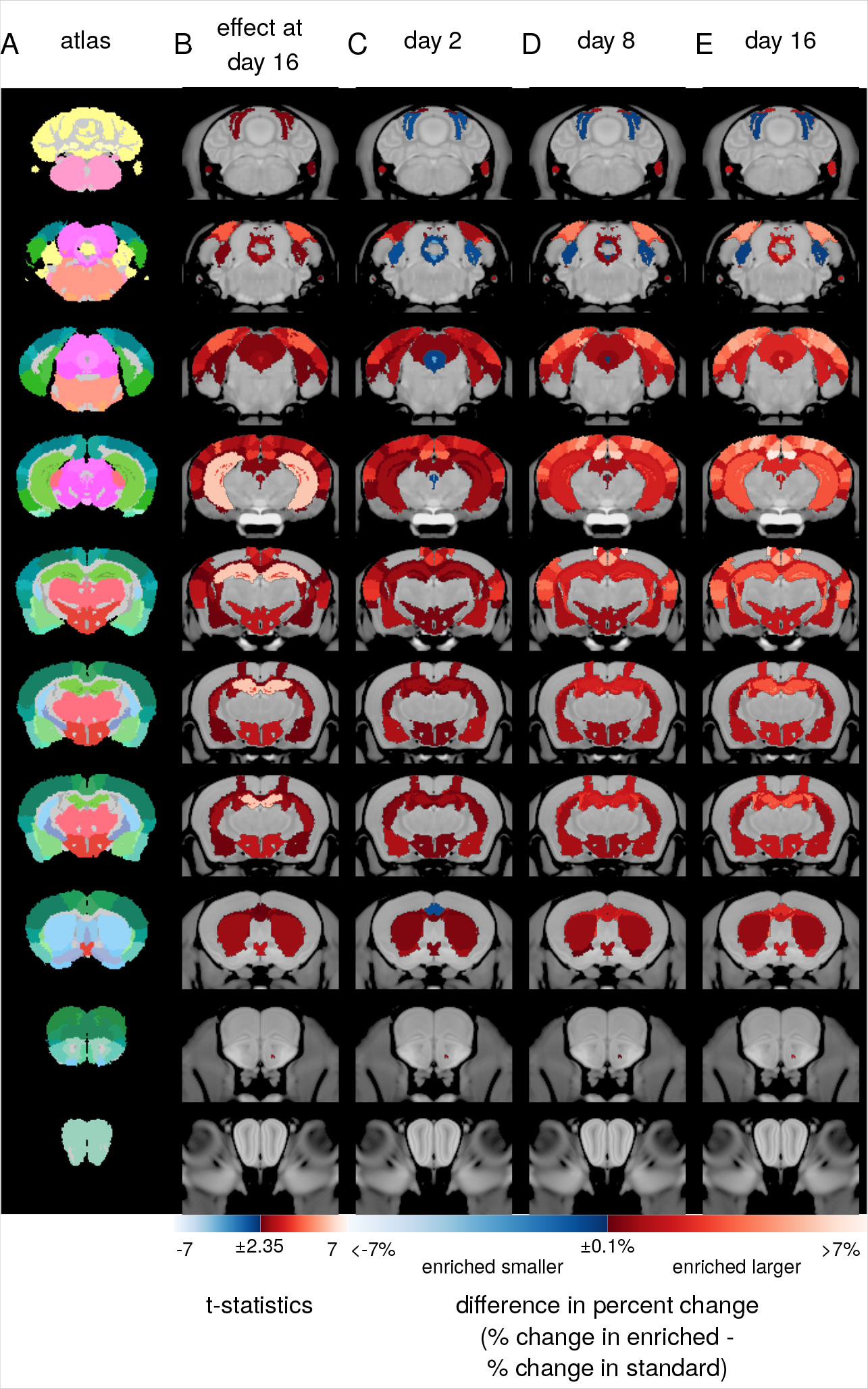
Environmental enrichment changes adult mouse neuroanatomy within 2 days and has progressively larger/more widespread effects over 16 days. Coronal images taken from the average anatomical magnetic resonance (MR) image of all mice in the study are shown overlaid with A) atlas labels delineating 159 regions in the brain; B) t-statistics (df=100) for the effect of enrichment from the linear model centred at day 16; and C-E) maps showing the difference in percent change between enriched (N=30) and standard housed (N=31) wild-type mice at day 2 (C), day 8 (D), and day 16 (E). Structures for which the growth rate was larger in enriched mice versus standard housed mice are shown in red, while structures for which the growth rate was smaller are shown in blue. Note that a larger growth rate in enriched mice can mean a relative reduction in volume loss. For example, after 16 days, there was a loss of cerebellar volume in both enriched and standard housed animals, but this effect was larger in standard housed mice. Only structures where there was a significant effect of enrichment at day 16 at a 10% false discovery rate (FDR) or less are shown.

**Figure 2:**
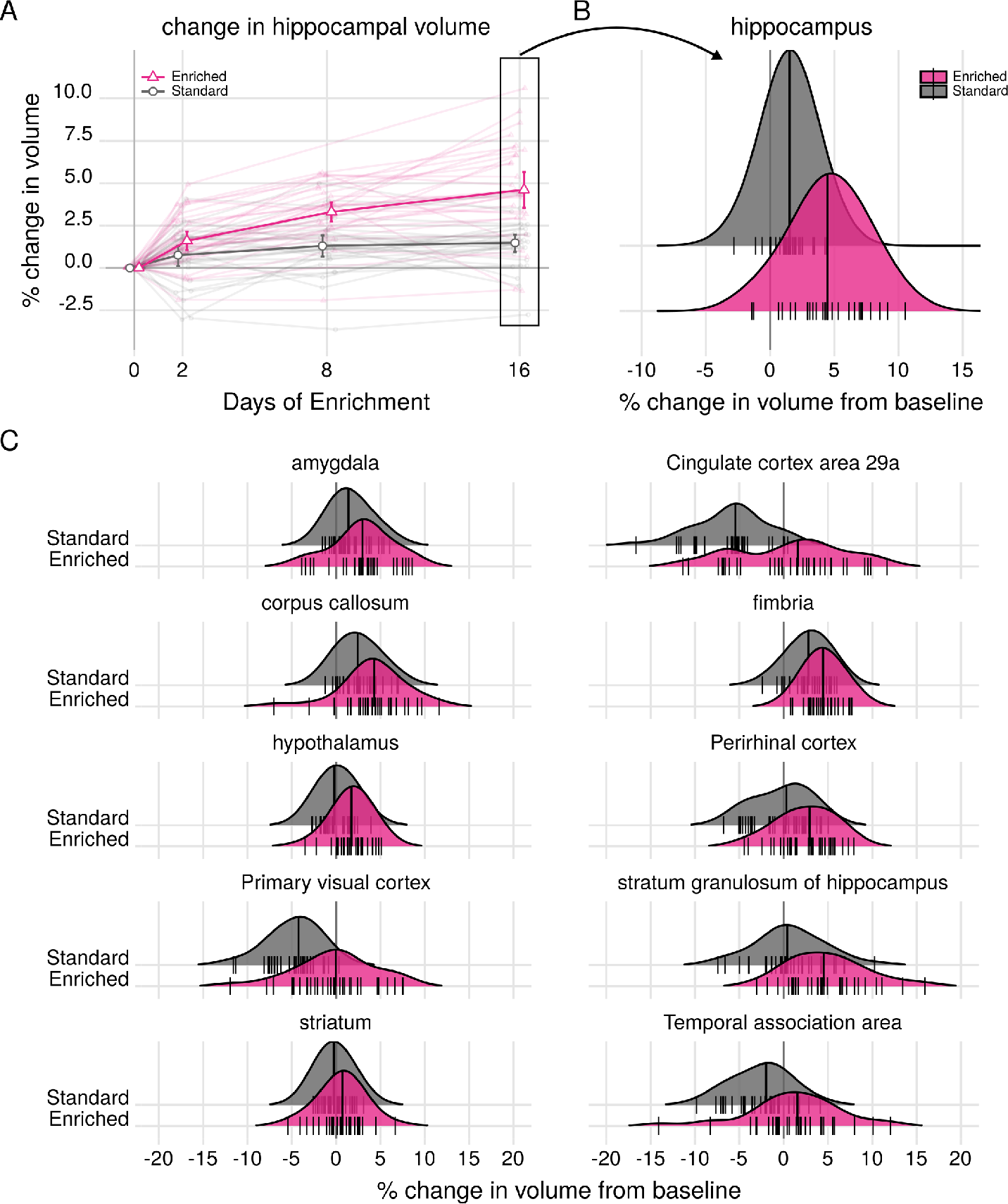
Sixteen days of enrichment leads to brain-wide changes in adult neuroanatomy. A) Percent change in hippocampal volume from baseline volume at the start of housing treatment: wildtype enriched (N=30, shown in pink) vs standard housed controls (N=31, shown in grey). Points are group means. Error bars are 95% confident intervals. Faded points and lines are individual mice. There was a significant effect of enrichment on the growth rate of the hippocampus (t(222)=6.93, q=6.76e-9). B) Density plot showing the distribution of changes in hippocampal volume after 16 days of standard or enriched housing. Sixteen days of environmental enrichment is associated with a significant increase in hippocampal volume (t(100)=6.03, q=2.52e-6), resulting in a mean 4.61% increase (sd=2.88%) in hippocampal volume between baseline and day 16 in enriched mice versus 1.48% increase (sd=1.48%) in standard housed animals. C) Sixteen days of enrichment is associated with a significant increase in volume in 39 structures (relative to standard housed wildtype mice). Density plots show the distribution of changes in volume in 10 representative structures from our atlas chosen to illustrate the range of effects. Small lines are individual subjects. Large line indicates group median.

The regions where the effect of 16 days of enrichment was the largest were the hippocampus (t(100)=6.03, q=2.52e-6), dentate gyrus (t(100)=6.00, q=2.52e-06), the primary visual cortex (t(100)=4.52, q=0.0007), and secondary visual cortex (lateral area, t(100)=4.53, q=0.0007). After 16 days, the hippocampus grew by 4.61% (sd=2.88%) in enriched mice compared to 1.48% (sd=1.48%) in standard housed mice. The effects of enrichment on hippocampal volume were rapid: the hippocampus grew by 1.60% (sd=1.58%) within 2 days in enriched mice versus 0.75% (sd=1.60%) in standard housed animals (t(100)=2.95,p=0.004).

After 8 days, there was a significant difference between enriched and standard housed mice in the volumes of 13 structures, including the hippocampus, dentate gyrus, and several subregions of the cingulate and visual cortices (q < 0.1 for all, see Supplementary Tables 6-8). After 16 days, significamt changes were observed in 39 structures, including the amygdala (t(100)=2.46, q=0.07); hippocampus (t(100)=6.03, q=2.52e-6), and dentate gyrus (t(100)=6.00, q=2.52e-06); white matter tracts such as the corpus callosum (t(100)=2.47, q=0.07), fimbria (t(100)=2.82, q=0.04), and cingulum (t(100)=3.68, q=0.02); the striatum (t(100)=1.89, q=0.04) and nucleus accumbens (t(100)=2.91, q=0.03); some lobules of the cerebellum (e.g. lobule 6 declive: t(100)=2.46, q=0.07); and the cingulate (t(100)=0.52 to 3.99, q=0.7 to 0.003), ectorhinal (t(100)=3.22, q=0.02), perirhinal (t(100)=3.52, q=0.01), somatosensory (hindlimb region, t(100)=2.55, q=0.07), visual (t(100)=2.77 - 4.52, q=0.03 to 0.0007 for 5 regions in our atlas), temporal association (t(100)=3.24, q=0.02), medial parietal association (t(100)=3.57, q=0.02), and primary auditory (t(100)=3.28, q=0.02) areas of the cortex (Figures 1, 2, S2, and S3, Supplementary Tables 6-8).

### 2.2 Effects of enrichment on brain anatomy are only partially mediated by exercise

We next asked whether we could dissociate the contribution of different components of environmental enrichment to brain wide changes in neuroanatomy. To test the hypothesis that the effects of enrichment on brain anatomy are mediated by increases in exercise, we placed an additional cohort of mice in cages containing exercise wheels and then imaged them at the same timepoints as described above. The distance run can be highly variable between mice [22] so the mice placed in exercise cages were housed singly, and odometers were attached to the wheels. To control for any effects of social isolation on brain anatomy, an additional cohort of mice was housed singly in standard cages (Isolated Standard group) and also imaged.

Previous studies show that both enrichment and exercise increase hippocampal neurogenesis, although they appear to do so via different cellular mechanisms [23, 24]. Consistent with this, exercise also increased the rate of growth of the hippocampus compared to mice housed in both isolated and group standard housing (Figure 3, S4, S5 interaction between exercise and day: t(352)=4.09 q=0.004 for comparison to standard housed mice and t(352)=3.89, q=0.019 for comparison to isolated standard). The effect of exercise on hippocampal volume was not significant at day 16 after correction for multiple comparisons (t(157)=2.87,p=0.005, q=0.18); however, the cumulative distance run after both 8 and 16 days significantly positively correlated with the change in hippocampal volume (Figure S6, t(36)=7.56, p=6.09e-09 and t=5.61(36), p=8.53e−7).

**Figure 3:**
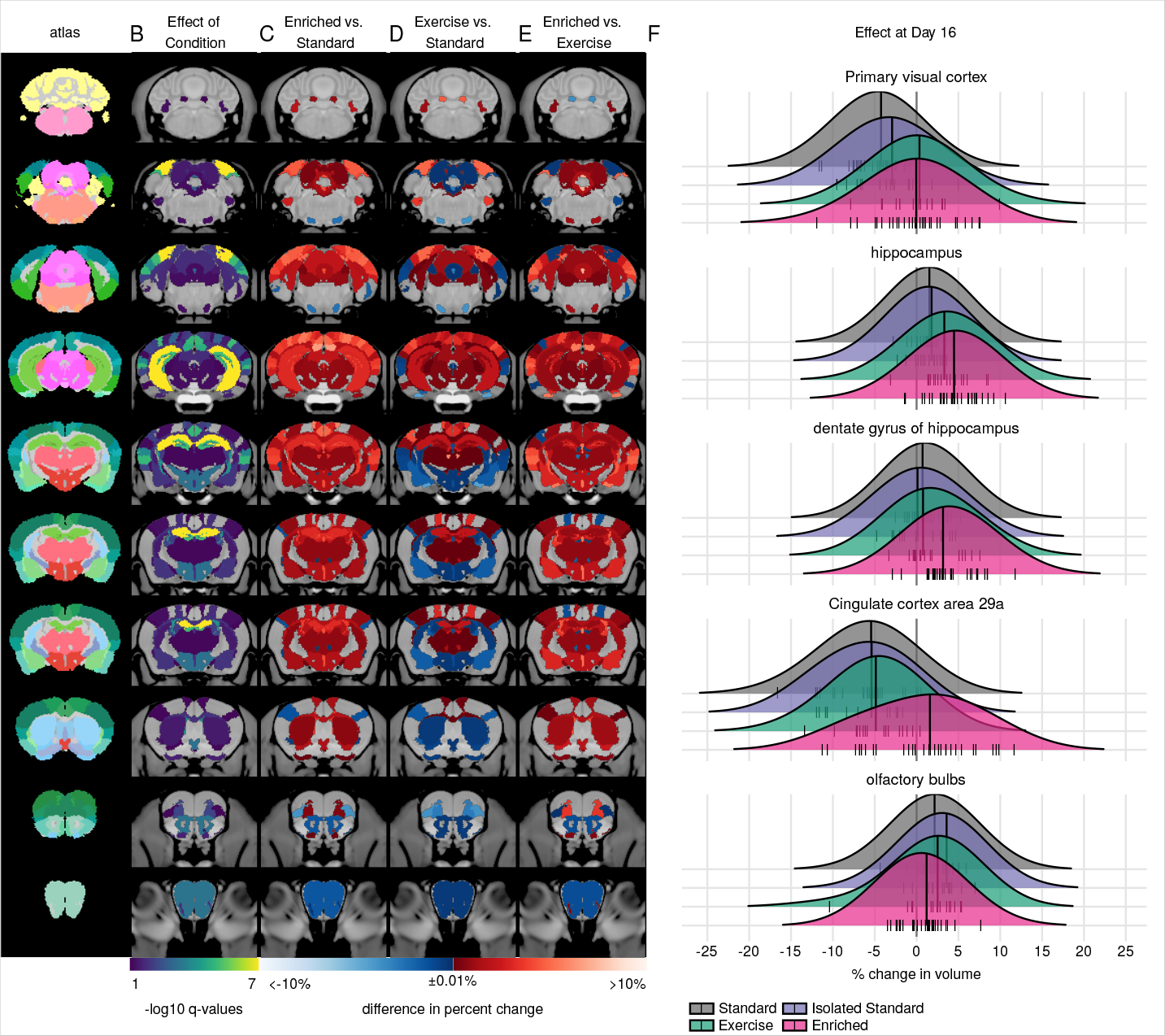
Effects of enrichment on brain anatomy are only partially mediated by exercise. Coronal images taken from the average anatomical MR image of all mice in the study are shown overlaid with A) atlas labels delineating 159 regions in the brain; B) negative log10 transformed q-values (FDR-adjusted p values) showing structures where there was a significant interaction between housing condition and day or significant effect of condition at day 16 of housing. A-log10 q value of 1 corresponds to q=0.1 while a-log10 q value of 7 corresponds to q=1e–7; and C-E) maps showing the difference in percent change between enriched (N=30) and standard housed (N=31) wildtype mice (C), exercise (N=19) and standard housed wildtype mice (D), and enriched and exercise wildtype mice (E) for each structure in our atlas. Structures for which the growth rate was larger in the enriched or exercise group versus the reference group (standard or exercise) are shown in red, while structures for which the growth rate was smaller are shown in blue. Only structures where there was a significant effect of housing condition at day 16 or a significant housing-day interaction at a 10% FDR are shown. (F) Density plots show the distribution of changes in volume in 4 regions of interest where there was a significant effect of housing treatment on structure volume after 16 days. Enrichment prevents a loss of volume in the primary visual cortex and cingulate cortex; increases the growth of the hippocampus and dentate gyrus; and decreases growth of the olfactory bulbs. Exercise alone increases hippocampal volume and prevents volume loss in the visual cortex. Small lines are individual subjects. Large line indicates group median.

Compared to exercise alone, enrichment had an even larger effect within the hippocampal formation and was associated with a larger growth rate in the dentate gyrus and stratum granulosum of the hippocampus (t(352)=2.78, q=0.046; t(352)=3.38, q=0.014). Outside the hippocampus, exercise recapitulated only a small portion of the effects of enrichment on the brain (Figure 3,S4,S5). There was a significant effect of exercise on the growth rate of 8/159 structures when compared to mice group-housed in standard cages, including an increase in the growth rate of the cingulate cortex area 29c (t(352)=3.09, q=0.049), primary visual cortex t(352)=5.42, q=1.79e–5), secondary auditory cortex t(352)=3.67, q=0.015), paraflocculus white matter (t(352)=3.32, q=0.032), and fasiculus retroflexus (t(352)=3.35, q=0.032), and a decrease in the growth rate of the superior olivary complex (t(352)=−3.02, q=0.055) and lateral orbital cortex (t(352)=−3.14, q=0.048, see Supplementary Tables 6-8). When the exercise cohort was directly compared to the animals housed in enriched cages, we found 23 structures in which enrichment altered the growth trajectory compared to those with access to an exercise wheel alone. Together, these data show that the effects of enrichment on brain anatomy cannot be primarily attributed to increased exercise.

### 2.3 Social interaction has few effects on brain anatomy

Interestingly, when the group standard housed mice were compared to those housed singly in standard cages, we found no brain regions with altered growth rates. After 16 days, only the flocculus white matter differed between standard housed and isolated standard groups and was significantly smaller in isolated animals compared to group housed (t(162)=−3.83, q=0.028).

Taken together, our results indicate that increases in exercise cannot explain the effects of enrichment on brain anatomy. Additionally, our data suggests that the effects of enrichment also cannot be ascribed to social interaction effects. While we cannot rule out the possibility that the quality or types of social interactions differ when mice are housed in enriched versus standard cages, environmental enrichment was associated with widespread cortical and subcortical growth, even though both enriched and standard cages contained the same number of mice. Given this, our results imply that the volumetric changes we observe following enrichment are most likely due to to cognitive and/or sensorimotor stimulation associated with learning to navigate a novel, complex environment and the episodic memory formation that results from this experience.

Given that the mesoscopic changes in neuroanatomy associated with enrichment appear to be driven by cognitive stimulation rather than exercise, we hypothesized that such changes may be attenuated by disrupting the animals’ ability to form long-term memories. In recent years, the molecular basis of memory formation has begun to be illuminated [25, 26]. One of the genes most strongly implicated in learning and memory is CREB [27–29]. CREB and its family of transcription factors have been shown to be critical for long-term memory formation in a number of species, including *Aplysia*, *Drosophila*, and mice. Disruptions to CREB are associated with impaired long-term memory on spatial learning tasks such as the Morris Water Maze [30, 31], impaired LTP [30], and unstable hippocampal place cells [32], suggesting an impaired ability to encode space. Moreover, CREB is a known regulator of many cellular processes stimulated by learning and environmental enrichment, including activity-dependent dendritic growth and cell survival [33, 34]. Given this, we next tested whether CREB is also required for the volumetric changes in neuroanatomy associated with environmental enrichment. To do so, we exposed mice with a targeted deletion of the two primary isoforms of CREB (CREB *alpha* and *delta*) to the same housing environments (enriched, group-housed standard, exercise, and isolated standard) and imaged them as described previously.

To ensure that wildtype and CREB-deficient mice experienced similar levels of enrichment, a subset of mice (3 cages, total N=15 mice) were observed for 3 hours after being placed into the enriched cages. All mice explored the cage, and within 3 hours, all mice regardless of genotype had reached the top level where the food was located. Thus, there does not appear to be an effect of CREB genotype on willingness to explore an enriched environment.

### 2.4 CREB*αδ* is not needed for changes in brain anatomy induced by environmental enrichment, exercise, or social interaction

Our primary hypothesis was that loss of CREB would attenuate the effects of environmental enrichment on brain anatomy. Surprisingly, there were only 4/159 structures in the brain where genotype modulated the effect of enrichment on growth rates (i.e. where there was an interaction between genotype, housing condition, and day of housing). The superior colliculus and cingulate cortex area 29a grew at a slower rate in enriched CREB*αδ*+/− mice versus in enriched wildtype mice (t(1017)=−3.37, q=0.048 for both, though this was not significant after 16 days), while the posterior commissure grew at a faster rate (t(1017)=3.33, q=0.048). Similarly, the quadratic component of growth was reduced in the paraflocculus of enriched CREB*αδ*−/− mice versus enriched wildtype mice (t(1017)=−3.45, q=0.09), but again this did not result in any differences at day 16. In fact, after 16 days of housing treatment, there were no voxels or structures where there was a significant interaction between genotype and housing condition (Figures 4, 5, and S7). Therefore CREB genotype does not modulate the effects of environmental enrichment on brain anatomy. Additionally, there were no structures in which genotype modulated the effect of 16 days of exercise or social interaction on structure volume.

**Figure 4:**
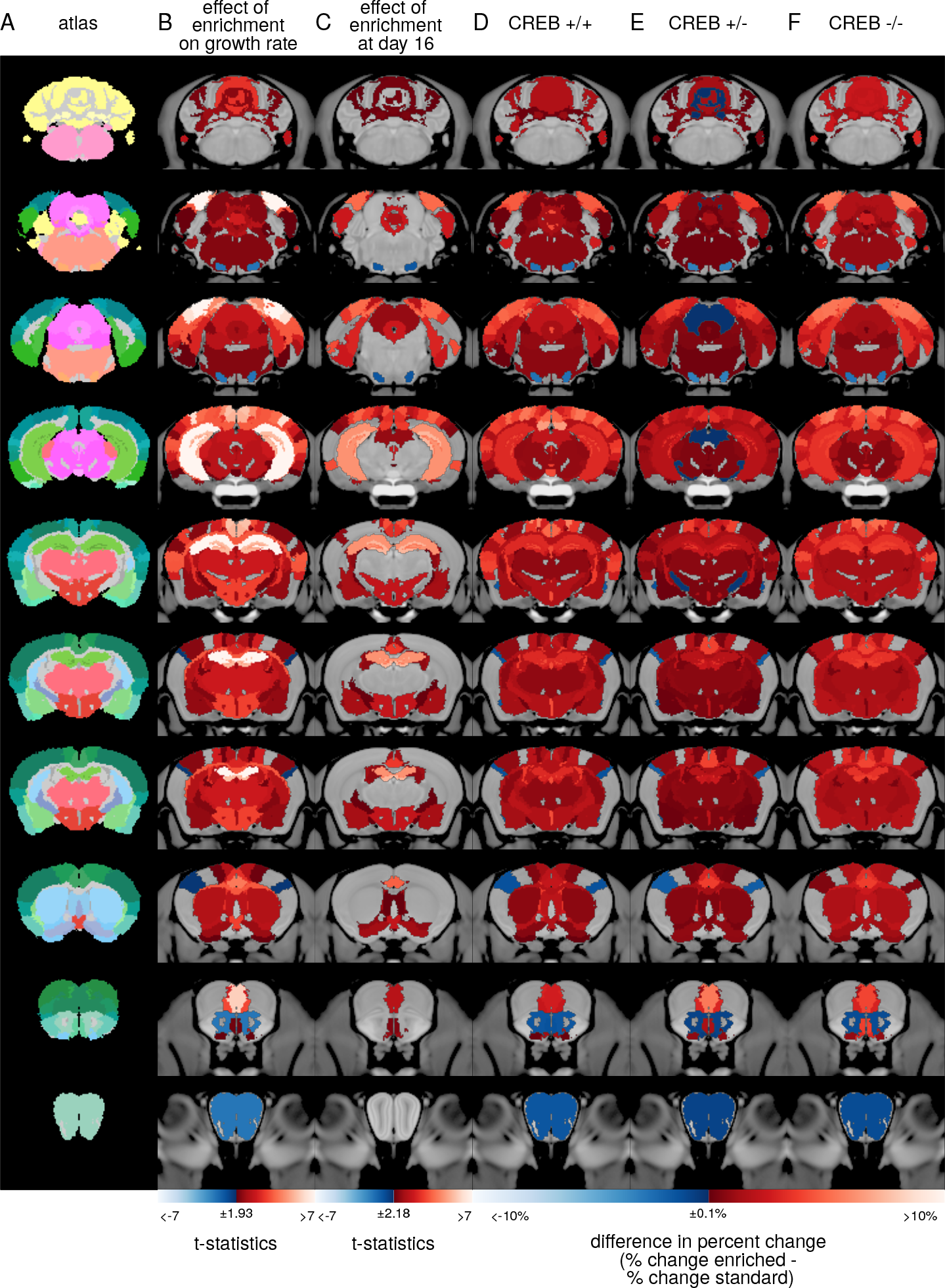
Loss of CREB does not affect volume changes induced by environmental enrichment. Coronal images taken from the average anatomical MR image of all mice in the study are shown overlaid with A) atlas labels which delineate 159 regions in the brain; B) t-statistics showing structures where there was a significant effect of enrichment on the linear component of growth rate (interaction between enrichment and day) for the model including all genotypes of mice; C) t-statistics showing a significant effect of enrichment at day 16 of housing; D-F) maps showing the difference in percent change between enriched and standard housed wildtype (CREB*αδ*+/+) mice (D), CREB*αδ*+/− mice (E), and CREB*αδ*−/− mice for each structure in our atlas. Sample sizes are shown in table S1. Only structures where there was a significant effect of housing condition at day 16 or a significant housing-day interaction at a 10% FDR are shown. There were no structures where there was a significant genotype-condition interaction after 16 days. Red colours indicate structures where enriched mice grew at a faster rate or had a larger volume after 16 days when compared to standard housed mice. Blue colours indicate structures that grew at a slower rate or had smaller volume after 16 days relative to standard housed mice.

**Figure 5:**
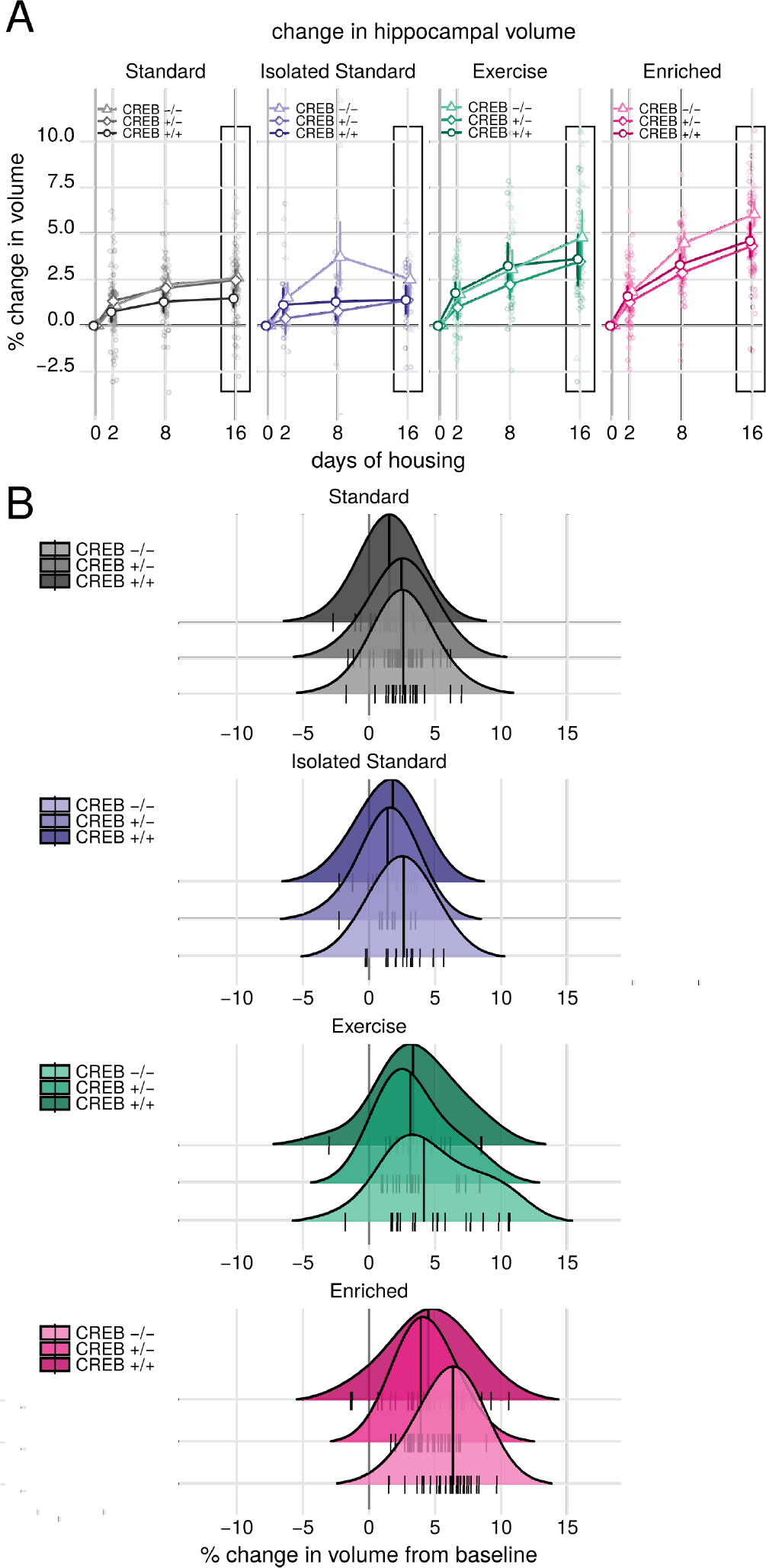
CREB is not needed for hippocampal volume increases associated with enrichment and exercise. A) Percent change in hippocampal volume from baseline over the course of 16 days of housing in an enriched environment (N=26-35/genotype), standard lab housing (N=26-34/genotype), social isolation (N=12-26/genotype), or a cage containing an exercise wheel (N=18- 22/genotype). Enrichment increases the growth rate of the hippocampus relative to standard housing (t=6.75, q=1e-8), but CREB genotype does not modulate this effect (condition-genotype-day interaction: χ^2^(18)=27.5,p=0.07 uncorrected; CREB*αδ*−/− x Enrichment x Day interaction: t(1017)=0.279, p=0.78). Solid points and lines are group means. Faded points are individual mice. Error bars are 95% confidence intervals. B) Density plots showing the distribution of changes in hippocampal volume in wildtype and CREB deficient mice after 16 days of housing treatment. After 16 days, there was a significant effect of housing condition on hippocampal volume (χ^2^(12)=182,p=2e-16) but no genotype-housing interaction (χ^2^(24)=30,p=0.18). Large lines are group medians. Small lines are individual mice.

These results suggest that contrary to our initial hypothesis, the neuroanatomical volume changes induced by enrichment do not require CREB-dependent transcription. Interestingly, an earlier study by Tsien et al. found that environmental enrichment increased synapse and dendritic spine density in CA1 of wildtype mice and in mice lacking NMDARs in this area [35]. NMDAR activation can lead to phosphorylation of CREB and activation of CREB-dependent signalling. Although the role of CREB was not investigated in that study, the finding that NMDAR-activity is not necessary for spinogenesis in the hippocampus suggests that CREB-dependent signalling may also be unnecessary. One hypothesis is that the volume changes reflect neuronal remodelling [13, 14]; however, it may be that neither NMDAR activity nor CREB are required for this plasticity.

### 2.5 Loss of CREB does impair memory formation, but does not block anatomical changes associated with training

A concern arising from these results was whether the ability of CREB-deficient mice to form episodic memories is actually intact. To confirm whether, as reported in previous studies, CREB deficient mice do have impaired memory formation, we trained a separate cohort of wildtype and CREB deficient mice on the spatial and non-spatial Morris Water Maze (MWM). Additionally, since we have previously found that spatial maze training is associated with an enlargement in hippocampal volume [13], we also asked whether loss of CREB would block the neuroanatomical changes associated with a more specifically cognitive, behaviourally-focused task. Accordingly, following water maze training, the brains of wildtype and CREB deficient mice were imaged with MRI. For comparison to our prior study [13], the brains were imaged 8 days after water maze training using ex-vivo (rather than *in-vivo*) MRI.

Mice of all genotypes improved in learning the location of the hidden platform in the spatial version of the MWM (Figure S8). However, CREB*αδ*−/− mice learned at a slower rate than CREB*αδ*+/− and CREB*αδ*+/+ mice (interaction between day and genotype: χ^2^(8)=65.268,p=4.27e–11, interaction between day and CREB*αδ*−/−: t=−5.53,p=3.5e–8, Figure S8). In addition, after 6 days of training the CREB*αδ*−/− mice travelled significantly further than wildtypes to locate the platform (t(168)=3.327, p=0.001). CREB*αδ*+/− mice also learned at a slightly slower rate (t)=−1.98,p=0.048) but did not differ significantly from wildtypes in their final performance (t(168)=−0.661,p=0.551).

CREB*αδ*−/− mice also displayed impaired spatial memory on a probe test performed 24 hours after the last training trial (Figure 6A). There was a significant interaction between genotype and pool zone on both the percent of the total path and the percent time spent in each quadrant (χ^2^(6)=36.10,p=2.64e-6 and χ^2^(6)=32.95,p=1.07e-5). CREB*αδ*+/+ and CREB*αδ*+/− mice spent significantly more time searching the quadrant of the pool in which the platform had been located during training (target vs. other quadrants: p < 0.001 for all comparisons). This memory for the platform location was impaired in CREB*αδ*−/− animals, who spent an equal amount of time searching the target and nearby left quadrant (t(292)=2.572,p=0.3009, Figure 6A) and significantly less time in the target quadrant than wildtype animals (t(292)=3.654, p=0.0158).

**Figure 6:**
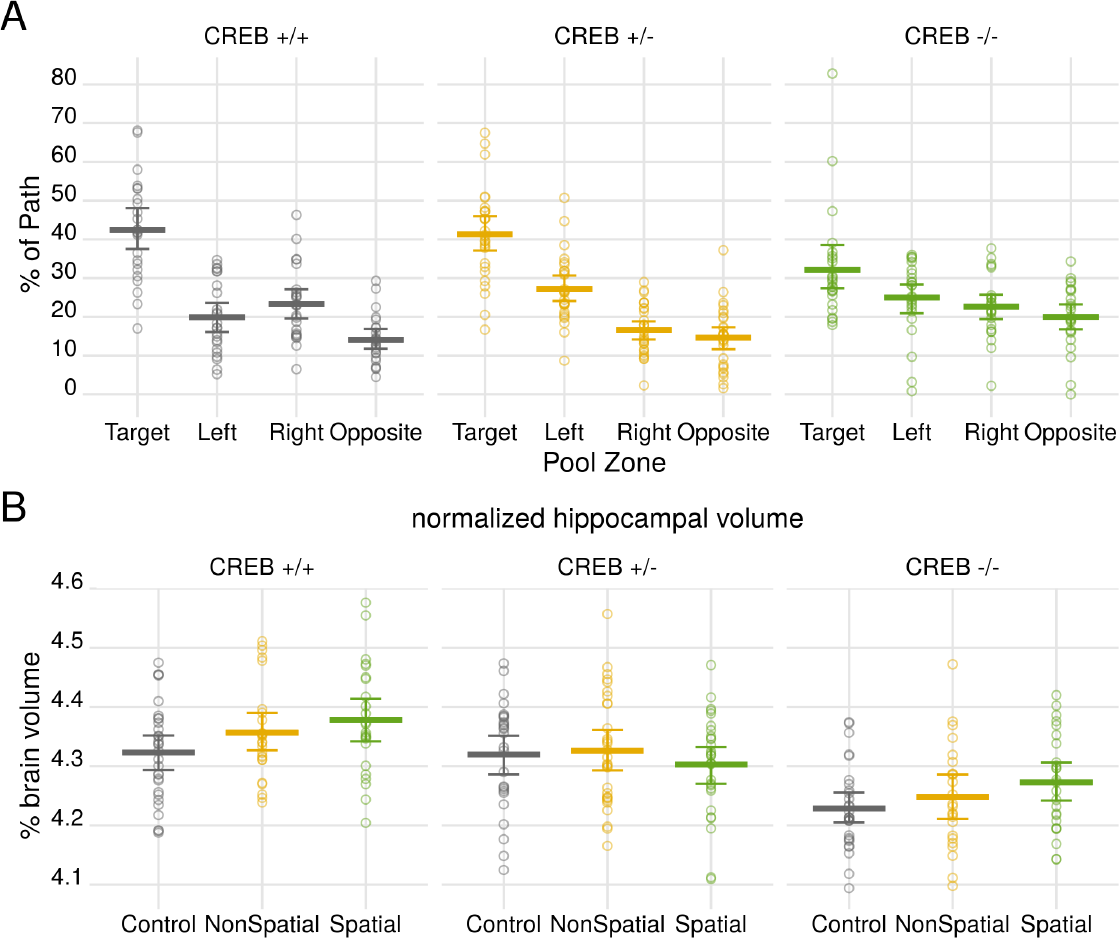
Loss of CREB impairs spatial memory formation but does not attenuate hippocampal volume changes associated with water maze training. A) There was a significant interaction between genotype and quadrant on the percent of the total path spent in each quadrant (χ^2^(6)=36.10,p=2.64e-6). CREB*αδ*+/+ and CREB*αδ*+/− mice show a strong preference for the target zone compared to non-target zones (target vs. other quadrants: p < 0.001 for all comparisons). CREB*αδ*−/− mice had a significantly reduced preference for searching in the target zone (t(292)=3.654, p=0.0158), indicating an impaired spatial memory. B) Effect of 6 days of maze training on normalized hippocampal volume. Normalized hippocampal volume is significantly larger in spatially trained mice compared to control animals (t(237)=2.276,p=0.025). There was no significant interaction between genotype and training condition on normalized hippocampal volume (F(4,233)=0.85,p=0.495). Lines are group means. Error bars are 95% confidence intervals. Points are individual mice.

To ensure loss of CREB impacts spatial learning specifically and to test for genotypic differences in motivation and/or locomotor activity, a separate group was trained on a non-spatial version of the water maze. CREB genotype only subtly altered the rate at which mice learned the non-spatial version of the MWM (Figure S8 interaction between day and genotype: χ^2^(8)=53.23 p=9.74e–09). CREB*αδ*−/− mice travelled further to the platform on day 1 (t=4.805,p=2.19e–06). However, for the remainder of training, there was no significant difference between the genotypes. Additionally, all animals trained on the non-spatial MWM searched the pool randomly during the probe test, showing no preference for a particular platform location (Figure S8).

Overall, the MWM experiments showed that CREB*αδ*−/− mice do retain some ability to learn but have impaired spatial memory. This is consistent with previous studies showing that loss of CREB leads to impairments in spatial memory and particularly that it impairs long-term but not short-term memory formation [29]. This data suggests that spatial memory within the context of environmental enrichment was effectively targeted by this model.

Since loss of CREB induced the predicted behavioural phenotype, we next examined the impact on the hippocampal volume changes associated with maze training. Because there was a significant effect of CREB genotype on overall brain volume (F(2,239)=18.36,p=3.84e-8) and animals were imaged only once ex-vivo, volumetric data here is presented normalized to total brain volume. Although the effect size is reduced compared to our previous study, hippocampal volume was significantly larger in spatially trained mice compared to control animals (Figure 6: Effect of condition (F(2,233)=3.03,p=0.050; spatial vs. control: t(237)=2.276,p=0.025). Moreover, there was no significant interaction between CREB genotype and training condition (F(4,233)=0.85,p=0.495), and hippocampal volume was larger in spatially trained CREB*αδ*−/− mice compared to control CREB*αδ*−/− mice (t=2.079,p=0.0411) when this genotype was analyzed separately. Thus loss of CREB does not attenuate the effects of spatial maze training on hippocampal volume.

In summary, our data show that environmental enrichment has rapid effects on brain anatomy that become larger and more widespread with prolonged enrichment. However, the effects of enrichment on brain anatomy cannot be attributed to social interaction or increased exercise alone. Instead, they likely arise from cognitive or sensorimotor stimulation and/or interactions between these factors and exercise/social interaction.

Moreover, we find that neuroanatomical volume changes induced by environmental enrichment do not require CREB. This means that although the effects of enrichment appear to be due to learning about a novel environment, the signalling pathway mediating these effects is distinct from the canonical CREB-dependent transcription pathway implicated in learning and memory formation.

One limitation of the mouse model used in this study is that it is not a full knockout of CREB since this mutation is perinatal lethal [36]. The most abundant isoforms of CREB are the *δ* and *α* isoforms; the *β* isoform is much less prevalent in the wildtype brain [37, 38]. In the wildtype brain, phosphorylated CREB heterodimerizes with other CREB proteins and with the CREB-family members activating transcription factor 1 (ATF-1) and cAMP-responsive modulator proteins (CREM) [39]. The heterodimer can then bind CRE sites in the genome and drive gene expression, with the help of additional transcriptional coactivators [39]. Targeted deletion of the *α* and *δ* isoforms results in a 90% reduction of CREB in the CREB*αδ*−/− mice and a 40% reduction in the heterozygotes; however, this genetic targeting also results in compensatory up-regulation of CREB*β* [37] and both the activator and repressor forms of CREM [40]. Consistent with this finding, we found roughly 2 fold higher total CREB mRNA expression in CREB*αδ*−/− mice and 1.5 fold higher total CREB mRNA in CREB*αδ*−/− mice compared to wildtype animals (Figure S9). As observed in prior reports, CREM was also upregulated in CREB*αδ*−/− mice but not in heterozygotes (Figure S9). The expression of ATF-1 and other related genes, including CREB binding protein and CREB-related transcription coactivator (Crtc1) was not affected by loss of CREB*αδ*.

Given this compensatory upregulation, it is possible that the CREB*αδ*−/− and CREB*αδ*+/− mice used here have sufficient residual CREB function to mediate any CREB-transcription-dependent structural changes. However, a study by Pandey et al. found that deletion of CREB*αδ* abolishes CREB and/or CREB/CREM DNA-binding [41]. Despite upregulation of CREB*β* and CREM, in the absence of CREB*αδ*, CRE-DNA binding is abolished in the amygdala, hippocampus, cortex, and cerebellum. This study therefore suggests that although CREB*β* and CREM are upregulated in CREB*αδ* deficient mice, there is still not CREB-dependent transcription in these animals, arguing against a role for residual CREB expression in our results.

More importantly, though, our data show that experience-dependent changes in brain structure volume do not require normal spatial learning and memory formation. Increases in hippocampal volume were still observed in animals with learning and memory deficits.

### 2.6 Enrichment and exercise induce distinct transcriptional pathways

Our data suggests that CREB-dependent transcription is not required for neuroanatomical plasticity. We next investigated other signalling pathways that might be driving these effects. We used RNA-sequencing to identify the transcripts differentially expressed in mice exposed to either an enriched environment or exercise wheel (versus standard housed controls). To determine whether the same transcriptional pathways were activated in wildtype and CREB*αδ* mutants, samples were taken from all genotypes of mice. Tissues were taken from regions of flash-frozen brains that showed a large volume change: the dorsal and ventral dentate gyrus and the occipital cortex.

Environmental enrichment was associated with significant (q < 0.05) upregulation of 887 genes and significant downregulation of 607 genes, relative to mice housed in a standard cage and covarying for transcriptional differences across regions. Within these genes, 151 were up or downregulated more than 20%, including genes involved in myelin-production (*Egr2*), neurogenesis (*S100a9*), immediate early genes (*fos*), as well as cytokines and related genes implicated in inflammation and immune response (*Ccr2*, *Lcn2*). In contrast, exercise was associated with upregulation of only 30 genes and downregulation of 19, of which 8 overlapped with the genes differentially expressed with enrichment (Figure 7). This data is consistent with our neuroimaging results showing that the effects of enrichment cannot be explained by exercise alone.

**Figure 7:**
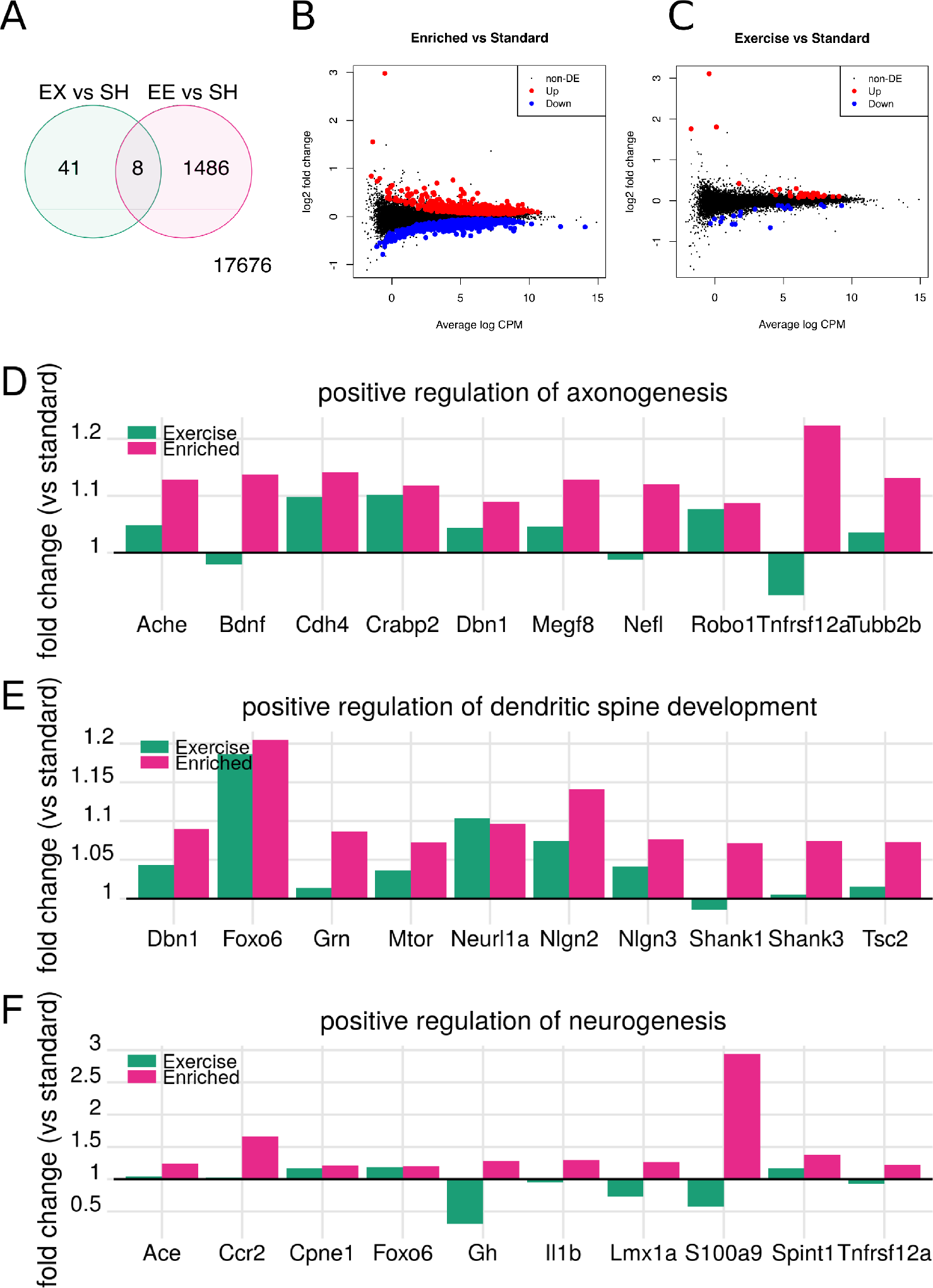
Environmental enrichment drives changes in transcription that cannot be explained by exercise alone. A) Venn diagram showing number of genes significantly differentially expressed in enriched (EE) or exercise (EX) versus standard housed (SH) mice after controlling for genotype and brain region variation. B) Mean difference (MD) plot showing the log2-fold change (log FC) and average abundance (log counts per million (CPM)) of each gene. Genes significantly differentially up- or down-regulated in EE vs SH mice are shown in red and blue, respectively. C) MD plot showing genes significantly differentially expressed in EX vs SH mice. Differential expression of genes associated with D) positive regulation of axonogenesis; E) positive regulation of dendritic spine development; and E) positive regulation of neurogenesis. Bar plots show fold change in gene expression in enriched (pink) or exercise (green) housed mice versus standard housed controls. Plots show mean fold-change across all regions in 10 genes associated with each gene ontology term. N=4-6 mice/genotype/condition/region.

Additionally, although there were 4595/19211 genes that were differentially expressed amongst the different genotypes of mice (CREB*αδ*−/−, CREB*αδ*+/−, and CREB*αδ*+/+), there was no evidence of genotype-enrichment interactions on gene expression. Again, consistent with our imaging data, this suggests that CREB genotype does not modulate the effects of enrichment on the brain.

### 2.7 Gene sets underlying neuroanatomical volume changes

Our next goal was to identify candidate gene sets and biological processes underlying the neuroanatomical volume changes seen with enrichment. As reviewed by Zatorre *et al*, there are several candidate processes through which experience-dependent volume changes may occur [15]. These include increases in neurogenesis; neuronal remodelling, including increases in dendritic spine density and axonal sprouting; gliogenesis and/or proliferation of microglia; angiogenesis; and changes to the extracellular matrix [15]. To determine whether genes associated with these processes were differentially expressed in mice exposed to enrichment or exercise (relative to standard housed mice), we used gene set enrichment analysis. Specifically, we investigated whether gene sets associated with the following gene ontology (GO) terms were differentially expressed with exercise or enrichment: positive regulation of neurogenesis; positive regulation of dendritic spine development; positive regulation of gliogenesis; positive regulation of angiogenesis; positive regulation of extracellular matrix organization; positive regulation of synapse structural plasticity; positive regulation of axonogenesis; positive regulation of microglia cell migration; and positive regulation of microglia activation.

In mice exposed to environmental enrichment, across the 3 brain regions, there was a significant upregulation of genes associated with positive regulation of axonogenesis (84 genes, q=0.001), dendritic spine development (57 genes, q=0.007), synapse structural plasticity (4 genes, q=0.013), and neurogenesis (470 genes, q=0.013) (Figure 7 and S10). Consistent with this, barcode plots show upregulation of genes that positively regulate these processes (Figure S10).

There was also significant downregulation of genes associated with positive regulation of microglia cell migration and activation (q=0.01 and q=0.003), though there were only 1-2 genes associated with each of these sets, respectively (Figure S10). While there was not evidence of directional changes (up or down-regulation) in gene expression across the remaining gene sets, we did find evidence of differential expression of genes associated with angiogenesis (118 genes, q=0.01 for differential expression, q=0.29 for up-regulation). Gene sets associated with positive regulation of extracellular matrix organization and gliogenesis were not differentially expressed (18 and 64 genes respectively, q=1 and 0.12).

To further investigate the biological processes and cellular components associated with enrichment, we used goseq to perform an unbiased test for enrichment of gene ontology categories within our set of differentially expressed genes [42]. We identified over 1000 GO terms over-represented within our list of genes differentially expressed with enrichment so we then used REVIGO to summarize the list. Amongst the top over-represented GO cellular components were extracellular region (p=0.001), cell (p=0.00001), neuron projection (p=10^−9^), and synapse (p=10^−10^) (Figure 8). The GO biological processes behaviour (p=0.01), cellular homeostasis (p=10^−7^), and signalling (p=0.001) were also significantly over-represented amongst our gene set. This data implies that neurogenesis and structural changes to the neuron may underlie the volume changes we see with MRI. Future studies could test this hypothesis using similar genetic targeting strategies as used here.

**Figure 8:**
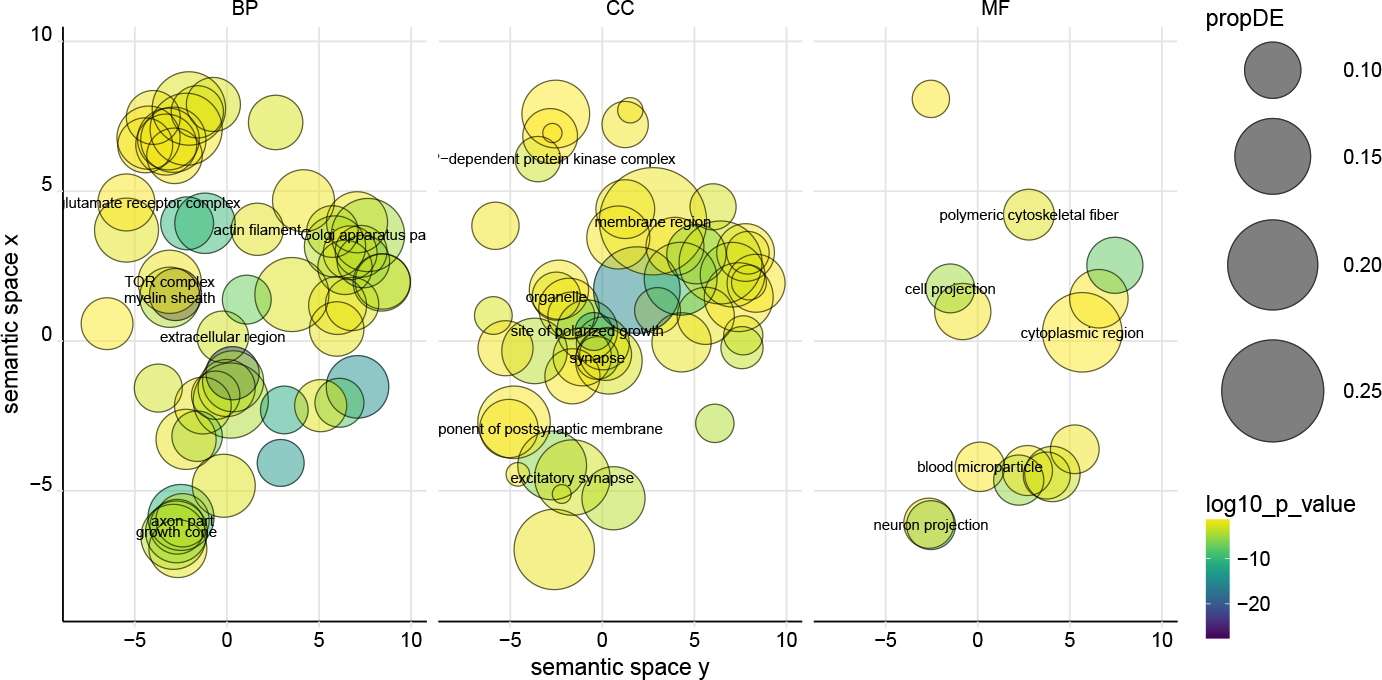
Gene ontology analysis with go-seq and REVIGO for genes differentially expressed in enriched mice versus standard housed controls. Differentially expressed genes were significantly associated with gene ontology terms shown, including terms associated with neuronal remodelling. MF=molecular function. CC=cellular component. BP=biological process. Size of bubble corresponds to the proportion of differentially expressed genes (propDE=number of differentially expressed genes/total genes associated with GO-term). Colour corresponds to the log10 p value. Bubbles are spaced in semantic space, where more semantically similar terms are closer together.

When we tested for whether these gene sets were differentially expressed with exercise, we likewise found significant upregulation of genes involved in dendritic spine development (q=0.036) and a trend towards upregulation of genes that positively regulate neurogenesis (q=0.060), though interestingly, we did not find evidence for differential expression of genes implicated in axonogenesis (q=0.36). Thus, although the genetic pathways stimulated by enrichment and exercise appear to differ, similar biological processes appear to be targeted. The difference in the extent of neuroanatomical changes may reflect the extent of differential expression.

## 3 Conclusions

The study of plasticity has historically been the domain of cellular neurobiology, which has limited the study of experience-dependent changes to only a few brain regions and to model organisms. Recently, though, studies from our lab and others have shown that learning and experience can change brain anatomy over a period of hours or days at the mesoscopic level of resolution accessible by MRI. These studies, including the study reported here, show that in-vivo, whole-brain imaging of brain plasticity associated with learning and experience is possible. The advantage of MRI is the ability to image the whole-brain non-invasively, and thus obtain a spatially-unbiased understanding of the effects of experience on the brain. The challenge, however, has been relating the changes observable with MRI to changes at the cellular and molecular levels.

Here, we used in-vivo mouse MRI and genome-wide transcription analysis to bridge the gap between cellular neurobiology and human neuroimaging studies of plasticity. We found that environmental enrichment drives differential expression of a large set of genes involved in regulating neuronal structure and number and leads to wide-spread changes in cortical and subcortical brain anatomy that emerge as soon as 2 days after the start of enrichment. This shows that in-vivo MRI is a powerful means of examining experience-dependent changes over the whole brain, and these changes can occur rapidly after the onset of a new experience. We also found that neither the transcriptional nor neuroanatomical changes associated with enrichment can be explained by increases in exercise or social interaction alone. Rather, they appear to be driven by the cognitive and/or sensorimotor aspects of this experience. Surprisingly, though, genetic targeting of a canonical transcriptional pathway that regulates learning and memory processes did not attenuate the neuroanatomical changes that occurred. This implies that the phenomena detected by MRI are not driven by the classical molecular learning pathways.

Within three regions that underwent large volume changes, we found that enrichment was associated with differential expression of genes that positively regulate axonogenesis, dendritic spine development, synapse structural plasticity, and neurogenesis. This suggests that the changes detected with MRI reflect many of the same processes historically measured using more cellular techniques. In addition, the transcriptional dataset acquired here provides a rich set of possible genetic targets to further probe the ontogeny of neuroanatomical plasticity.

## 4 Methods

### 4.1 Mice

All studies and procedures were approved by The Centre for Phenogenomics (TCP) Animal Care Committee in accordance with recommendations of the Canadian Council on Animal Care, the requirements under the Animals for Research Act, RSO 1980, and TCP Committee Policies and Guidelines. We used adult male and female wildtype mice and mice with a targeted deletion of two CREB isoforms (B6/129 CREB*αδ* mutants: CREB*αδ*+/−, CREB*αδ*−/−) [30, 40]. Experimental mice were the F1 hybrid offspring of a cross of CREB*αδ*+/− mice on a 129 background with CREB*αδ*+/− mice on a C57BL6/J background. Wildtype mice are designated CREB*αδ*+/+. Prior to weaning, mice were ear-notched with individual markings to distinguish cage mates, and the ear punches were used for genotyping. Genotyping was performed using qPCR (Transnetyx). Mice were weaned at postnatal day 21 and then housed in 3–5 mice/cage. Males and females were housed separately. When possible, at least one mouse of each genotype from a given litter was included in the cage, and littermates of the same sex and genotype were assigned to different experimental groups. When litters did not contain one pup of each genotype and sex, litters born within 2–3 days were pooled so that cages contained at least one mouse of each genotype. Mice were maintained under controlled conditions (25 degrees C, 12/12 hour light/dark cycle, lights on at 7am) at the TCP in individually ventilated, sterile cages. Mice were provided a standard irradiated chow from Harlan (Harlan, Teklad Global 18% Protein Rodent Diet, 2018) and sterile water ad libitum.

The total number of mice used in this study is listed in supplementary tables S1 and S2. The sample sizes used for imaging experiments were determined prior to the study using power analyses. We generated simulated datasets and computed the sample size needed to detect a significant genotype-condition or genotype-condition-day interaction, assuming a 3% change in wildtype hippocampal volume and no change in hippocampal volume in CREB *αδ*−/− animals. The effect size of 3% was chosen as this was the effect size reported in a previous study from our lab [13].

### 4.2 Environmental enrichment and exercise treatment

Mice were randomly assigned to be placed in group enriched environment housing (Enriched), group standard housing (Standard), individual exercise-only housing (Exercise), or individual standard housing (Isolated Standard, N=12–35 per genotype and housing condition, see supplementary table S1). Mice were placed into their housing treatment at 8-9 weeks of age along with the rest of their cage mates (where applicable). The enriched environment was a Double Decker Rat IVC Green Line cage (Techniplast, Italy) with 3 levels of tubes and tunnels, nesting material, and a running wheel [19]. Each enriched environment cage was paired with a standard housing cage containing the same number of mice (3–5). The exercise-only cages contained only bedding, nesting material, and a running wheel with an odometer attached (5 inch diameter aluminium activity wheel, Lafayette Instruments, Lafayette, IN, USA). The distance each mouse ran was recorded continuously by the odometers and a log was taken each week day between 8–11 am. To control for the effects of social isolation, an additional group was placed in individual (isolated) standard housing. The mice in the isolated standard housing group were in identical cages to the exercise group but did not have running wheels.

### 4.3 *In vivo* magnetic resonance imaging

To obtain a reliable baseline measurement of neuroanatomy, we imaged the brains of mice using *in vivo* manganese-enhanced MRI three and four days prior to housing treatment. Mice were imaged 7 at a time on a multichannel 7T Varian scanner in individual saddle coils using a 3D gradient echo sequence (scan parameters: 90 *μ*m isotropic resolution, repetition time (TR)=29ms, echo time (TE)=5.37ms, flip angle=37°, matrix size=224 × 224 × 854, averages=5, 1 hour 40 min. scan time). Throughout the scan, bore temperature was maintained at 29°C, and 1-2% isoflurane was used to keep the mice anaesthetized. Respiration was monitored throughout the scan.

The mice were then imaged 2, 8, and 16 days after onset of housing treatment. Mice were imaged in the same coil across all timepoints. With the exception of the baseline scans, mice were injected with 50 mg/kg MnCl_2_ 24 hours before imaging. For the baseline scans, mice were injected only once 18 hours prior to the first scan. This was done to avoid toxicity due to repeat injections within 24 hours, since MnCl_2_ has a long half life.

To correct for small geometric distortions resulting from imaging in coils not in the centre of the magnetic field, coil-specific MR images of precision-machined phantoms were registered to a computed tomography (CT) image of the same phantom. The resulting distortion correcting transformations were then applied to all acquired images in a coil-specific manner.

### 4.4 Manganese chloride preparation

A 300 mM stock solution of MnCl_2_ was made by dissolving manganese chloride tetrahydrate (Sigma Aldrich 13446–34–9) in hydroclone purified water. For intraperitoneal injections, this solution was diluted in sterile saline to yield a 30 mM working solution. The animals were weighed prior to injection and given an appropriate volume of 30 mM stock to provide a 50 mg/kg dose of MnCl_2_ based on the animal’s weight. Previous studies from our lab indicate this dose does not have an adverse affect on behaviour [43].

### 4.5 *In vivo* image registration

An automated image registration pipeline was used to align all the *in vivo* MR images as previously described [20, 44–46]. Images were excluded from mice who died prior to the day 2 scan.

The registration was performed in two stages: first, for each mouse, the images from each timepoint were aligned to create subject-specific averages; and secondly, these averages were aligned to create a consensus average of all the images (i.e., all subjects and timepoints) in the study [9, 19, 47] (Figure S11a). For both the within-subject and between-subject alignment, the registration procedure includes both an affine registration and a series of non-affine (non-linear) registrations. The affine registration linearly aligns all the input images using a series of global rotations, translations, scales, and shears and relies on the mni autoreg tools [48]. To precisely align the images, the linearly aligned images are then locally deformed through an iterative nonlinear process using Advanced Normalization Tools (ANTS) [49]. The outputs of this image processing pipeline are 1) a consensus average representing the average neuroanatomy of all mice in the study; 2) an average image for each mouse (i.e. the average image over all time points); and 3) deformation fields relating each individual image to both the subject-specific and group averages.

The statistical analysis was performed using the log Jacobian determinants of these deformation fields as the dependent variable. The Jacobian determinants encode how much larger or smaller each voxel is than the subject-specific or group average [44, 45]. Two sets of Jacobian determinants were used: the first set of Jacobian determinants are referred to as ‘first level Jacobians’ (Figure S11b). These are computed using within-subject deformation fields and encode how the images of one individual from one time point differ from that individual’s average scan. The second set of Jacobians is computed using the deformation fields relating the subject-specific averages to the study-average (Figure S11c). These Jacobians represent how an individual mouse’s average scan differs from the group average. Together, these Jacobians encode how each time point for each mouse relates to the study average. The analysis of voxel-wise growth rates was performed using the first set of Jacobians, which represents within-subject changes and can be used to detect differences in growth rates. The analysis comparing the Jacobians at particular time points was done using the full Jacobians (within-mouse + between mouse Jacobians combined) relating each image to the study average.

To obtain volumes of different anatomical structures, all the linearly aligned images were automatically segmented using the MAGeT (multiple automatically generated templates) algorithm [50]. We used a pre-existing mouse brain atlas that segments the brain into 159 structures for this process [51–53]. Briefly, MAGeT generates a set of template atlases by non-linearly aligning the pre-existing atlas to a subset of the input images. Then, the input images are aligned to each of these templates. Finally, a voxel-voting procedure is used to choose the best label for each voxel [50]. This process produces an automatically generated, individual atlas for each input image, which allows us to compute the volume of each of the 159 segmented structures for each image. Of note, the hippocampal formation is segmented into three regions in our atlas: hippocampus (including CA1-3 and the subiculum); dentate gyrus; and separately, the granular cell layer of the dentate gyrus.

### 4.6 Statistical analysis of *in vivo* imaging data

All analysis was performed using the R statistical language (R Core Team, 2016, https://www.R-project.org) using the RMINC and lme4 packages (https://github.com/Mouse-Imaging-Centre/RMINC and [54, 55]). To determine how the volume of each voxel in the brain changed over the course of housing treatment, separate polynomial models were fit at each voxel testing for linear, quadratic, and cubic effects of days of housing on the log-transformed Jacobians. A similar analysis was performed for each of the 159 structures defined in our atlas. Linear mixed effects models were used with fixed effects of days of housing and random intercepts for each mouse. Mixed effects models are appropriate for longitudinal studies in which data from the same subjects are acquired at multiple timepoints, and so the datapoints are not fully independent. Moreover, compared to methods such as repeated measures ANOVAs, mixed effects models are better suited to handling missing data-points and/or imbalanced study designs.

The general model formula for the volume of a given structure or of the log Jacobian of a given voxel (*y*) for mouse *i* on day *j* is:

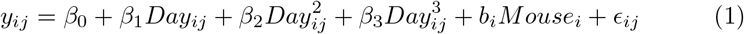

To determine the significance of adding/removing a term to the model, a second model was fit at each voxel omitting the terms of interest, and the models were compared using likelihood ratio tests using the Satterthwaite approximation for degrees of freedom [55]. The total number of scans used in a given model varied between 292 and 1338. As an example, to compare a model with a cubic effect of day to a simpler quadratic model, the quadratic model was also fit at each voxel:

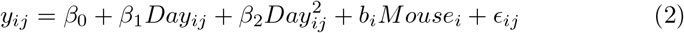

A test statistic *D* can then be computed from the maximum likelihood of the full model *L*_*f*_ (1) and the partial model *L*_*p*_ (2):

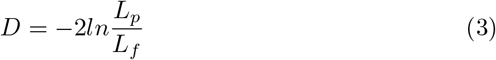

According to Wilks’ theorem [55], *D* follows a χ^2^ distribution with degrees of freedom equal to the difference between the number of parameters in the full model and that in the partial model. Using this approximation, we can compute p-values to determine the significance of the effect of interest.

There was a cubic effect on a 42% of the voxels whose volume was affected by day of housing (Figure S1) so a third-order (cubic) polynomial model was chosen to model the effect of day of housing. This analysis was then repeated to examine whether sex, genotype, housing treatment, or the interactions between these factors altered the trajectory of daily voxel growth. To do so, these terms were added as covariates. In other words the *β*_1_, *β*_2_, and *β*_3_ coefficients were allowed to vary with housing treatment, genotype, and/or sex. As described above, the models were compared using likelihood ratio tests to determine the significance of adding/removing a term to the model. T-statistics were also computed for each coefficient of interest. The resulting statistical maps were thresholded to control for multiple comparisons using a False Discovery Rate (FDR) threshold of q=0.10 [56]. This threshold was chosen as simulation studies from our lab show that at this threshold, we can recover volume differences of 10% with a false positive rate of less than 0.007% [57].

To determine the effect of housing, sex, and/or genotype at a particular day *d*, we used age-centered models. For each of the time points of interest (day 2, 8, and 16), a separate model was fit translating the growth effects such that they were 0 at each of the days of interest:

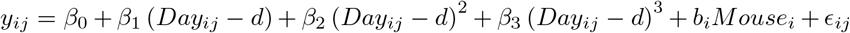

For simplicity, the model above is shown without interaction effects between day and other factors (e.g. housing condition), but interactions between the centered-day term and housing treatment, sex, and genotype were also included. This analysis was performed both at each voxel and for each structure.

In addition, for each mouse and at each voxel and structure, we computed the percent change in voxel/structure volume from baseline to each of our days of interest (i.e. percent change from day 0 to day 8). The Jacobian values and structure volumes obtained from the two baseline scans were averaged for this purpose. We then averaged these percent change maps to compute the mean percent change in voxel/structure volume for each genotype and housing condition at each time point.

### 4.7 Morris Water Maze

A second group of adult CREB*αδ* mice were trained on either the spatial (spatial MWM) or cued non-spatial (non-spatial MWM) version of the Morris Water Maze (see Table S2). Mice were randomly assigned to spatial, non-spatial, or no (control) training. Prior to water maze training, all mice were handled for 2 min/day for 5 days. Control mice were handled but not trained. For water maze training, mice learned to locate a hidden platform that was submerged in a 121 cm diameter pool of water made opaque using non-toxic white paint. The temperature of the pool was maintained at 25 degrees Celsius. A 12 cm diameter platform was submerged 5–8 mm below the surface of the water. For the spatial version of the maze, the platform location varied for each mouse but was constant over all trials, and distal cues were placed on the walls surrounding the pool to provide cues for navigation. For the non-spatial version of the maze, a curtain surrounded the pool which blocked the distal spatial cues from view. Lights were placed symmetrically around the top of the curtain to ensure the lighting level was similar in the spatial and non-spatial configurations of the task. In the non-spatial task, the platform location varied between 1 of 4 locations for each trial, and a red rectangular marker was placed on the platform to mark the platform location.

Mice were 12 weeks old at the start of training and were trained each day for 6 days, with a probe test on the 7th day. Each day, each mouse performed 2 blocks of 3 trials. The inter-trial interval was 15 seconds, and the inter-block interval was 1 hour. Before the first trial on day 1 only, each mouse was placed on the platform for 15 seconds. For each training trial, one mouse was taken from its cage and carried to the pool in the experimenter’s hands. The other mice remained hidden from view. The mouse was lowered into the pool with its nose facing the pool wall at one of 4 start locations (north, south, east, west). The start locations were randomly selected, but were the same for all mice of a given experimental group and always included the same number of locations close to versus far from the platform location. Immediately after placing the mouse in the pool, the experimenter went behind a curtain to be hidden from view and started the trial. Mice were given 60 seconds to find the platform location. The trial ended if the mouse found the platform, after which the mouse was left on the platform for 15 seconds before being returned to a recovery cage on a heating pad. If the mouse did not find the platform within 60 seconds, it was guided to the platform with the experimenter’s hands and left there for 15 seconds. 24 hours after the last trial on day 6, a probe test of memory was performed during which the platform was removed, and the mouse was allowed to search for the platform for 60 seconds. Both the spatial and non-spatially trained mice received the same probe test with the curtain open. All training was performed between 8am and 4pm by the same experimenter who was blinded to genotype. Video tracking of the mice was performed using the HVS Image Water 2020 Software using a tracking camera mounted to the ceiling above the centre of the pool.

### 4.8 Statistical analysis of behavioral data

For each trial, the following metrics were automatically recorded by the Water 2020 software: latency to reach the target platform (seconds), distance travelled to platform (metres), thigmotaxis time (seconds and % of total time), swim speed (metres/second), time spent in each pool quadrant (seconds and % of total time), and time spent in a circular zone (20 cm radius) surrounding the target (or equivalent zone in non-target quadrants). Behavioural analysis was performed in R using the lmerTest and lsmeans packages [54, 55, 58, 59]. Separate linear mixed effects models were used to determine whether there were main effects or interactions between day of training, genotype, and/or sex on each behavioural metric. Fixed effects of day of training, genotype, and/or sex were included along with random intercepts for each mouse. P-values were obtained by comparing the full model with the effect in question against the model without that effect using likelihood ratio tests. T-statistics and associated p-values for the parameter estimates were calculated using Satterthwaite’s approximations for degrees of freedom. A 4th order natural spline fit was found to be the best fit for the distance travelled data.

### 4.9 Brain preparation for *ex vivo* MRI

Mice were perfusion-fixed 8 days after MWM training as previously described [20, 45, 60]. Briefly, mice were first perfused via the left ventricle using 30 mL of room-temperature (25°C) phosphate-buffered saline (PBS, pH 7.4), 2 mM ProHance (gadoteridol, Bracco Diagnostics Inc., Princeton, NJ), and 1 *μ*L/mL heparin (1000 USP units/mL, Sandoz Canada Inc., Boucherville, QC) at a rate of approximately 1 mL/minute. Then, 30 mL of 4% paraformaldehyde (PFA) in PBS containing 2 mM ProHance was infused at the same rate. After fixation, the heads, skin, ears, and lower jaw were removed and the skull was allowed to postfix in 4% PFA at 4 degrees C for 24 hours. The samples were then placed in a solution of PBS, 2 mM ProHance, and 0.02% sodium azide (sodium trinitride, Fisher Scientific, Nepean, ON) and stored at 4 degrees C until imaging.

### 4.10 *Ex-vivo* MRI

Anatomical whole-brain images were acquired 16 at a time using a multi-channel 7.0-T Varian scanner and custom-built 16-coil solenoid array (Varian Inc., Palo Alto, CA) [45, 61]. Brains were imaged using a T2-weighted rapid acquisition with relaxation enhancement (RARE) scan at 40 *μ*m isotropic resolution. The MRI parameters used were: TR=350ms,TE=15ms, effective echo time=30ms, echo train length=6, matrix size=504×504×630, ∼14 hour scan time [62]. Images were distortion corrected as described above.

### 4.11 *Ex-vivo* image registration and structure volume computation

*Ex-vivo* MR images were aligned using the same automated image registration as described in section 4.5 except the within-subject registration was excluded since there was only one scan per subject. Likewise, structure volumes were obtained using the MAGeT (multiple automatically generated templates) algorithm with a pre-existing mouse brain that segmented the brain into 159 distinct regions [50–53].

### 4.12 Statistical analysis of *ex vivo* imaging data

All analysis was performed using the R statistical language (R Core Team, 2016, https://www.R-project.org) using the RMINC package (https://github.com/Mouse-Imaging-Centre/RMINC/). Linear models were used to determine the effects of genotype, training condition, or the interaction between these two factors on hippocampal volume. Sex was included as a covariate as it was determined that there was no interaction between sex and the other factors.

### 4.13 Preparation of brain for RNA sequencing

A subset of mice exposed to an enriched, exercise, or standard housing environment were used for RNA sequencing (N=4-6 per genotype per condition, see supplementary table S3). To limit confounding transcriptional variability, only male mice were used for RNA sequencing. Four days after the last scan, mice were sacrificed via cervical dislocation and the brains were extracted, flash frozen in 2-methylbutane and then stored at −80°C. For the dorsal hippocampus, brains were sliced coronally using a cryomicrotome at 200 *μ*m until reaching bregma −2.30. The brains were then removed from mounting position, rotated and remounted to the mounting position for horizontal slicing of ventral hippocampus tissue. Horizontal sections were sliced from interaural 3.24 mm to 0.92 mm. A 350 *μ*m diameter puncher was used to punch visual cortex, dorsal and ventral dentate gyrus region separately. Tissues were collected in cold 1.5 mL eppendorf tubes and stored at −80°C.

### 4.14 RNA sequencing

RNA extraction was performed using Qiagen RNeasy Micro kit (Qiagen, Cat 74004) with on-column DNase I treatment to remove genomic DNA contamination. The RNA was examined by Bioanalyzer 2100 (Agilent technologies, Santa Clara, USA). The RNA libraries were prepared in lab using Illumina TruSeq stranded total RNA LT set (CA RS-122-2301, Illumina Canada Ulc.) RNAseq data was collected with paired-end 100 bp reads of length using HiSeq 4000 at a depth of 25 M sequencing at McGill University and the Genome Quebec Innovation Centre.

### 4.15 Analysis of RNA sequencing data

RNA sequencing reads were checked for quality with the FastQC package and were aligned to the mouse mm10 genome with the STAR aligner. Counts of reads to annotated mm10 genes were also computed with STAR.

All differential expression analyses were done with the Bioconductor software using the edgeR and goseq packages [42, 63, 64]. We only retained genes with counts per million > 1 in at least 3 samples. To eliminate bias in library composition, normalization factors were calculated for each sample using trimmed mean of M values (TMM). Differentially expressed genes were identified using generalized linear models testing for main effects of housing condition and/or housing condition x genotype or region interactions. Specifically, we tested the following null hypotheses:

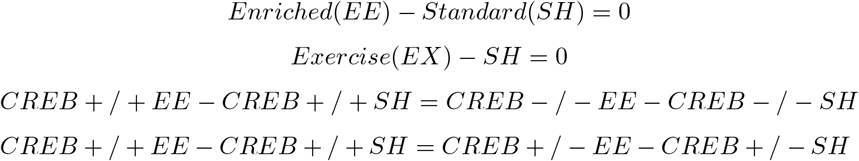

Multiple comparisons were controlled using a false discovery rate of q < 0.05.

Gene set enrichment analysis was performed using the rotation gene set tests implementation in limma for multiple gene sets (fry)[65–67]. First, we identified genes associated with the following gene ontology terms: 
- GO:0050769: positive regulation of neurogenesis
- GO:0060999: positive regulation of dendritic spine development
- GO:0014015: positive regulation of gliogenesis
- GO:0045766: positive regulation of angiogenesis
- GO:1903055: positive regulation of extracellular matrix organization
- GO:0051835: positive regulation of synapse structural plasticity
- GO:0050772: positive regulation of axonogenesis
- GO:1904141: positive regulation of microglia cell migration
- GO:0014008: positive regulation of microglia activation

Then we tested whether these sets of genes were significantly up-regulated in enriched or exercise versus standard housed mice. P values were adjusted using a FDR threshold of q < 0.05.

Unbiased gene ontology analysis was performed using goseq [42]. The resulting list of over-represented GO terms was summarized using REVIGO [68].

### 4.16 qRTPCR validation of RNAseq

The RNAseq data were validated with qRTPCR on 15 selected genes across the three brain regions (dorsal dentate gyrus, ventral dentate gyrus and visual cortex). cDNA conversion was done using 0.5 g RNA and Maxima First Strand cDNA synthesis kit for RT-qPCR (Cat K1642, Thermo Fisher Scientific Inc.). Probe based quantitative real time reverse transcription polymerase chain reaction (qRTPCR) for CREB1 and mouse Beta-2 microglobulin (B2M) gene transcripts was performed using a real time thermocycler (Lightcycler 480, Roche Applied Science). Sybr green based regular qRTPCR was performed for the remaining genes. The mouse B2M gene, a house keeping gene, was subjected to PCR amplification using B2M primer set without probe to control for equal loading. Forward and reverse primers and probes are specified in supplementary table S4. A delta-delta Ct method was used to calculate the relative fold gene expression between groups. Log fold change comparison was analyzed between RNAseq and qRTPCR.

The results show a high correlation between RNAseq and qRTPCR (CREB +/+: enrichment vs standard, r = 0.764, p = 0.001; CREB +/+: exercise vs standard, r = 0.896, p<0.0001; CREB +/−: enrichment vs standard, r = 0.894, p<0.0001); CREB +/−: exercise vs standard, r = 0.890, p<0.0001 (Figure S12 and supplementary table S5).

### 4.17 Data availability

MRI and behavioural data will be publicly released on Brain-CODE (http://braininstitute.ca/research-data-sharing/brain-code). All data is available by contacting Dulcie Vousden or Jason Lerch.

### 4.18 Code availability

The code for image registration (https://github.com/Mouse-Imaging-Centre/pydpiper) and statistical analysis (https://github.com/Mouse-Imaging-Centre/RMINC) is freely available online.

## 5 Acknowledgements

The authors gratefully acknowledge the assistance of the following people: Chris Hammill and Darren Fernandes provided advice on statistical analysis. Matthijs van Eede provided computing support. Ramy Ayoub and Lindsay Cahill assisted with exercise cage assembly. Jan Scholz and Jun Dazai designed the enrichment cage, along with RAG. Jeff Emmanuel assisted with pilot water maze experiments. Adelaide Yiu gave advice on behavioural training. The staff of TCP, including Vanessa Ashthrope and Kyle Duffin, assisted with breeding, genotyping, and housing of mice. Mark Henkelman and Derek van der Kooy provided stimulating discussion as this data was acquired. Yohan Yee assisted with Figure S11.

This study was supported by funding from an Ontario Graduate Scholarship, the University of Toronto, and the Hospital for Sick Children (D.A.V), the Canadian Institute for Health Research (J.P.L), the Hope for Depression Research Foundation (M.J.M), and the Ludmer Centre for Neuroinformatics and Mental Health Foundation (T.Y.Z).

## 6 Author Contributions

DAV designed the study, did the behavioural training, MR imaging, imaging and behavioural analysis, sample preparation for MRI and RNA seq, RNA seq analysis, and wrote the paper. AF did in-vivo MR imaging. LQ assisted with the RNA sequencing preparation and in-vivo MR imaging. SAJ and PWF contributed mice and assisted with study design and interpretation. MM oversaw the RNA sequencing. XW performed RNA extraction, library prep and qRTPCR validation. NO did RNA seq analysis. BD contributed to the image registration software. JD organized and purchased materials for RNA sequencing. LSN and BJN optimized pulse sequences for MRI. RAG assisted with study design. TYZ oversaw and designed the RNA sequencing experiments, was responsible for RNA seq and qRTPCR technical support and analysis, and wrote the RNA sequencing methods and results. JPL designed the study and oversaw the project and analysis. All authors contributed to revisions of the paper.

## 7 Conflict of Interest Statement

The authors declare no conflicts of interest.

## 10 Supplementary Figures and Tables

### 10.2 Supplementary Figure Captions

**Figure S1:**
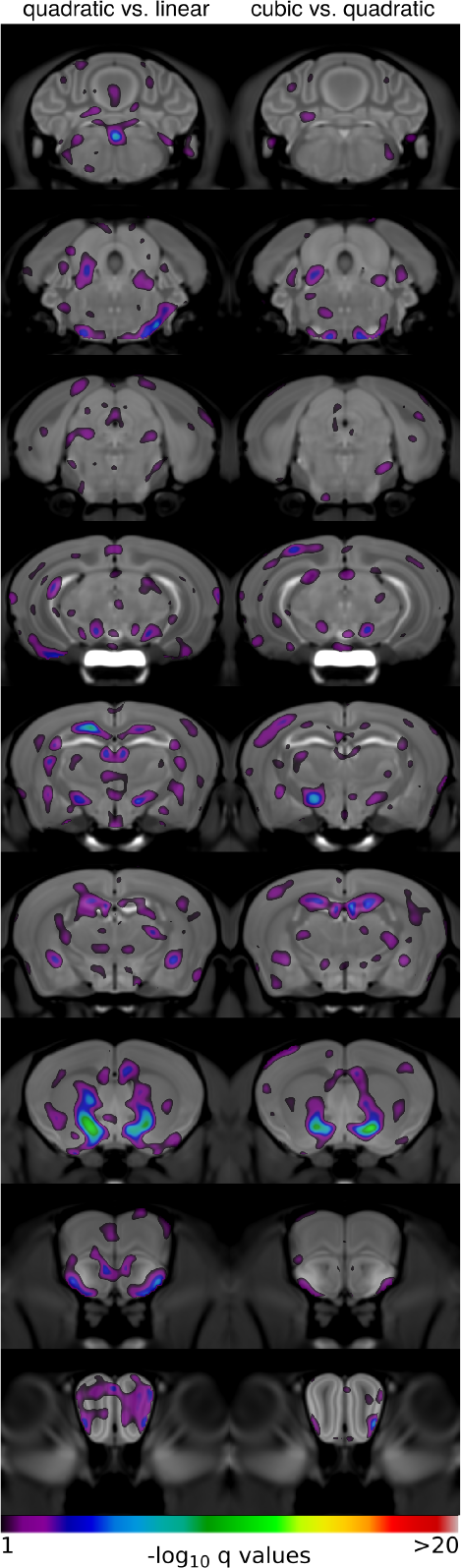
Comparison of linear, quadratic, and cubic effects of day on voxel volume. Polynomial models with a linear, quadratic, and cubic term for day of housing were fit at each voxel and compared with likelihood ratio tests. Coronal sections from the average anatomical MR image of all mice in the study are shown overlaid with FDR-corrected p values (q-values) for the likelihood ratio tests comparing models. Q-values are shown on a −log10 scale, thresholded at 10% FDR. Colours depict areas that were significantly better explained by a quadratic versus linear model (left) or cubic versus quadratic model (right). A −log10 value of 1 corresponds to an FDR corrected p value < 0.1.

**Figure S2:**
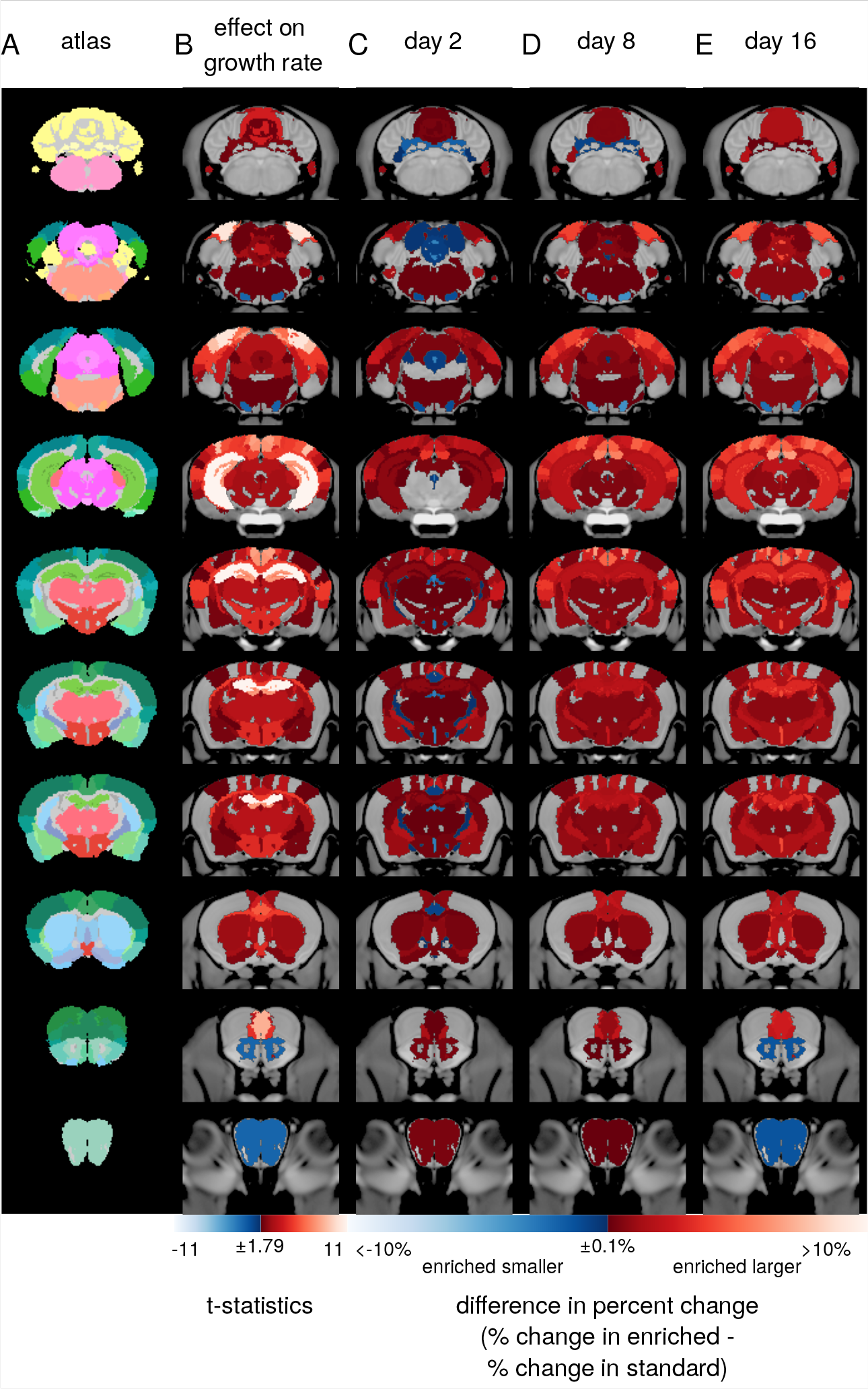
Environmental enrichment increases the growth rate of cortical and subcortical brain regions. Coronal images taken from the average anatomical MR image of all mice in the study are shown overlaid with A) atlas labels delineating 159 regions; B) t-statistics for the difference in the linear component of growth rate between enriched (N=30) and standard housed (N=31) wildtype mice; and C-E) maps showing the difference in percent change between enriched and standard housed mice at day 2 (C), day 8 (D), and day 16 (E). Only structures where there was a significant interaction between housing condition and day at a 10% FDR are shown. Note that this figure is comparable to figure 1 except that the interaction between enrichment and day of housing is shown versus the main effect of enrichment at day 16.

**Figure S3:**
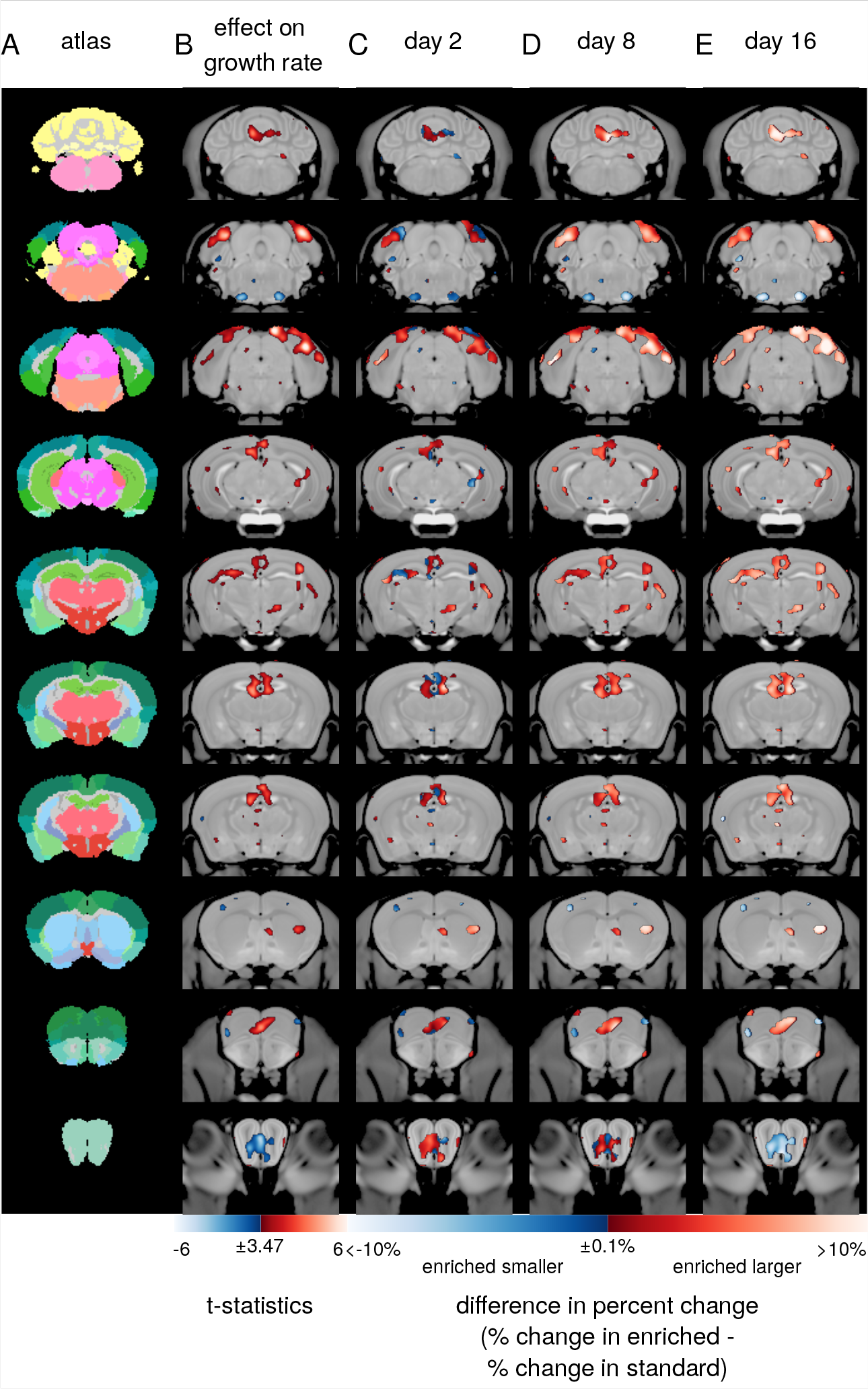
Voxelwise analysis of the effects of environmental enrichment on adult brain anatomy. Coronal images taken from the average anatomical MR image of all mice in the study are shown overlaid with A) atlas labels which delineate the brain into 159 regions; B) t-statistics for the difference in the linear component of growth rate between enriched (N=30) and standard housed (N=31) mice at each voxel; and C-E) maps showing the difference in percent change between enriched and standard housed mice at day 2 (C), day 8 (D), and day 16 (E) at each voxel. Only voxels where there was a significant interaction between housing condition and day at a 10% FDR are shown.

**Figure S4:**
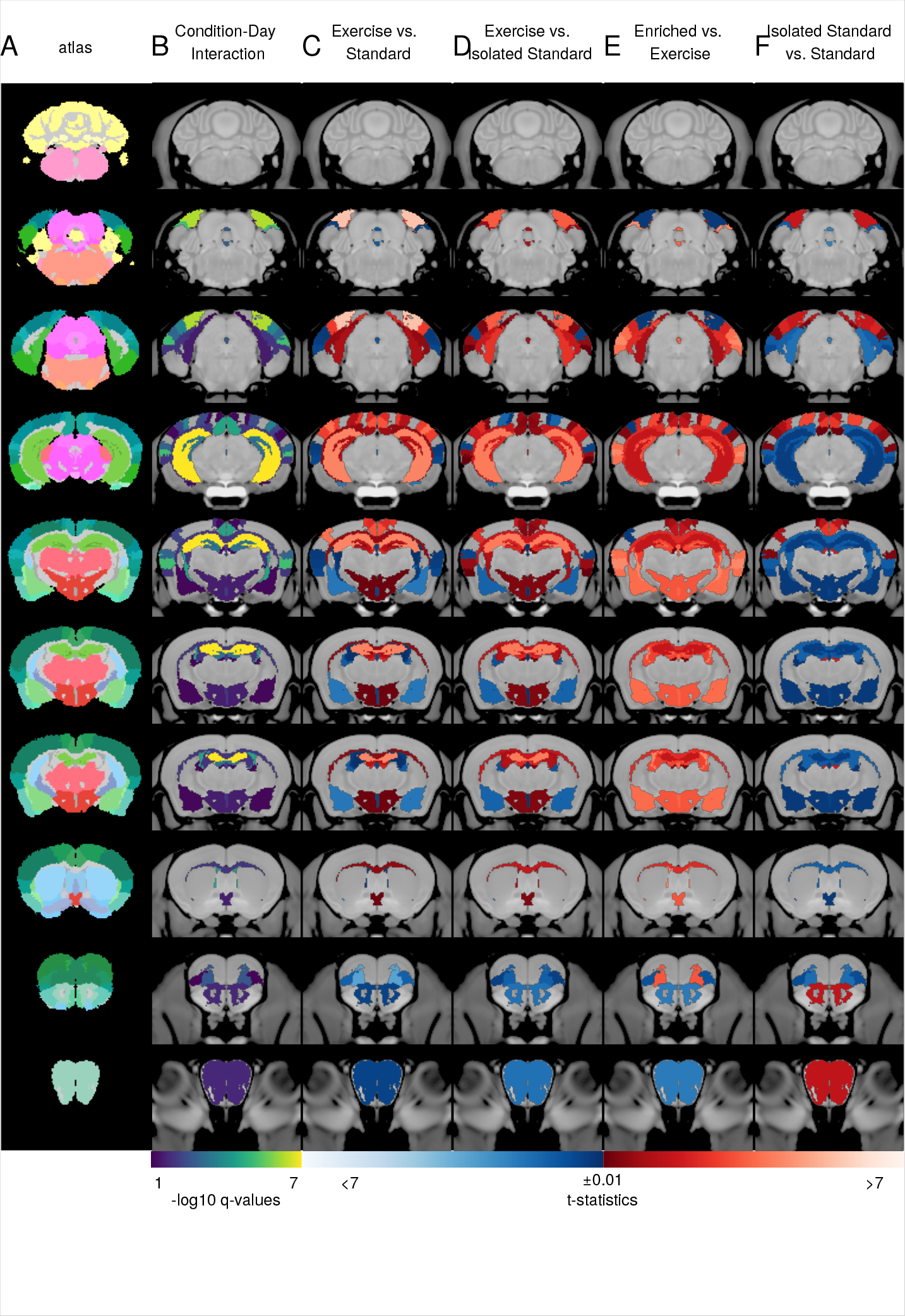
Effects of enrichment on brain anatomy are only partially mediated by exercise. Coronal images taken from the average anatomical MR image of all mice in the study are shown overlaid with A) atlas labels which delineates 159 regions in the brain; B) negative log10 transformed q-values (FDR-adjusted p values) showing structures where there was a significant interaction between housing condition and day. A −log10 q value of 1 corresponds to q=0.1 while a −log10 q value of 7 corresponds to q=1e–7; and C-E) t-statistics showing the difference in growth rate (condition-day interaction) change between enriched (N=30) and standard housed (N=31) wildtype mice (C), exercise (N=19) and standard housed wildtype mice (D), enriched and exercise wildtype mice (E) and isolated standard (N=17) versus standard housed mice (F) for each structure in our atlas. Structures in red had a larger growth rate in the first listed group versus the reference group while structures in blue had a smaller growth rate. T-statistics were masked using the −log10 q values shown in panel B. This means that t-statistics for all structures where there was a significant housing condition-day interaction at a 10% FDR are shown, even if the t-statistics were not significant. This is why, for example, t-statistics are shown for the effect of isolated standard housing on the growth rate of the hippocampus, even though this effect was not significant.

**Figure S5:**
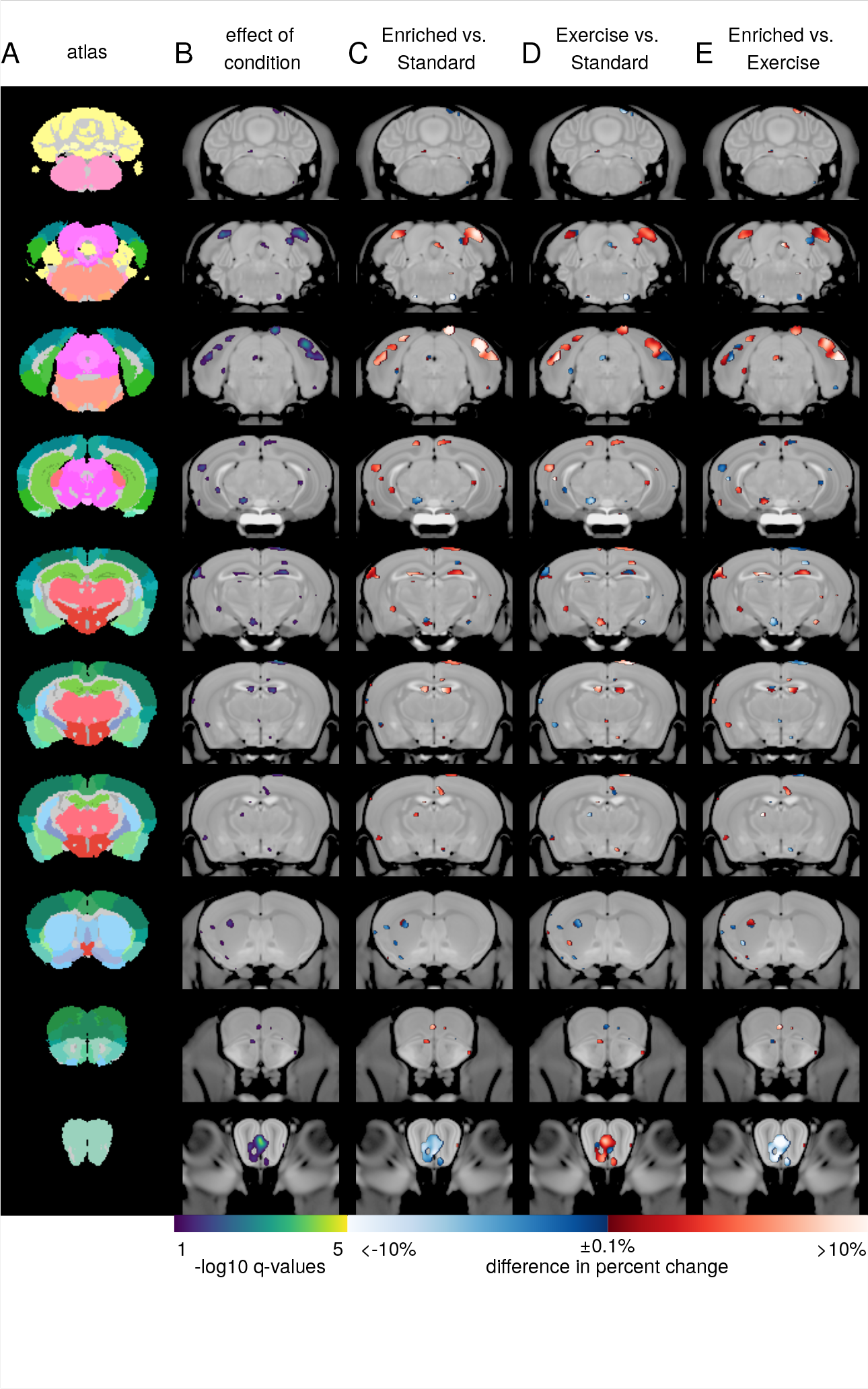
Voxel-wise analysis showing that the effects of enrichment on brain anatomy are only partially mediated by exercise. Coronal images taken from the average anatomical MR image of all mice in the study are shown overlaid with A) atlas labels which delineates 159 regions in the brain; B) Negative log10 transformed q-values (FDR-adjusted p values) showing structures where there was a significant interaction between housing condition and day or significant effect of condition at day 16 of housing. A −log10 q value of 1 corresponds to q=0.1 while a −log10 q value of 7 corresponds to q=1e–7; and C-E) maps showing the difference in percent change between enriched (N=30) and standard housed (N=31) wildtype mice (C), exercise (N=19) and standard housed wildtype mice (D), and enriched and exercise wildtype mice (E) at each voxel. Structures in red had a larger growth rate in the enriched or exercise group versus the reference group (standard or exercise) while structures in blue had a smaller growth rate. Only voxels where there was a significant effect of housing condition at day 16 or a significant housing-day interaction at a 10% FDR are shown. This represents the same analysis as shown in Figure 3 shown voxel-wise.

**Figure S6:**
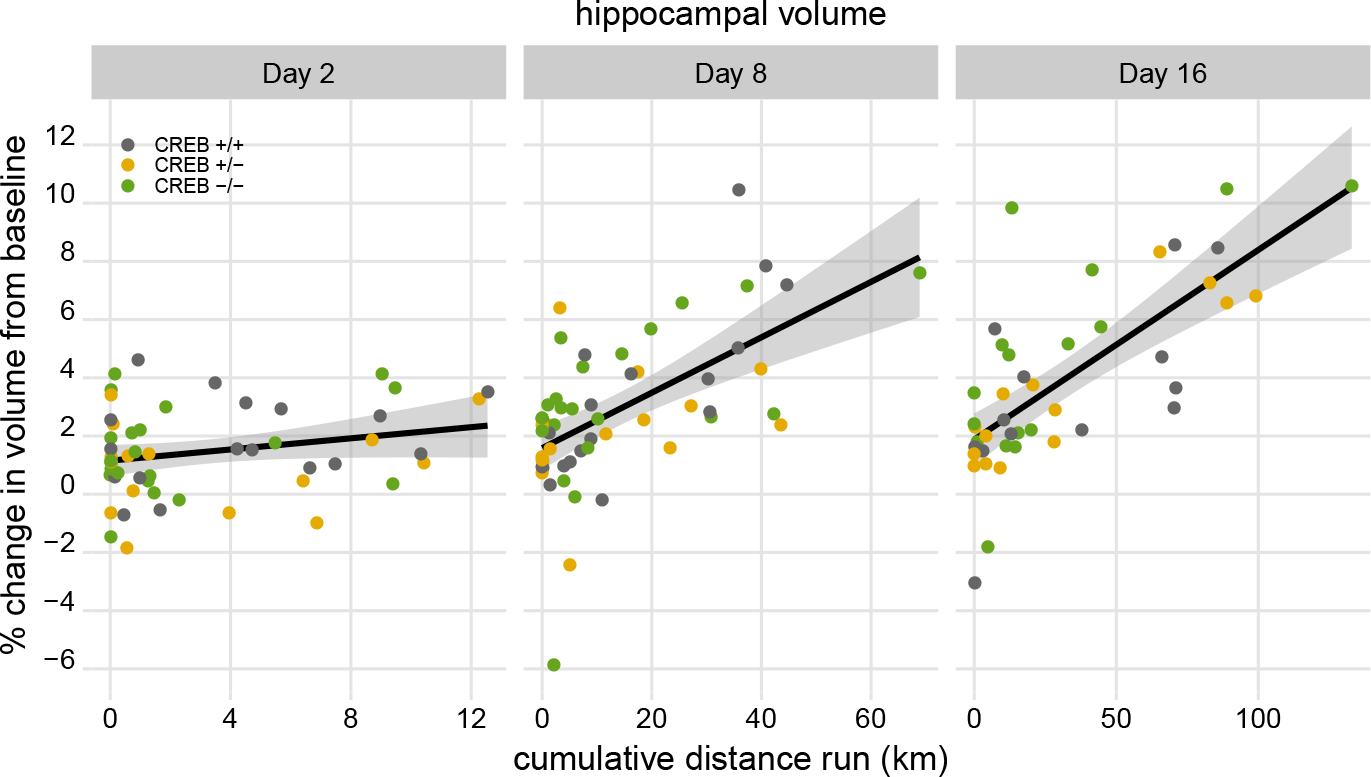
Hippocampal volume correlates with distance run. Figure shows correlation between cumulative distance run by mice housed in cages in exercise wheels and change in hippocampal volume after 2, 8, and 16 days of housing. There was a significant correlation between cumulative distance run and change in hippocampal volume after 8 and 16 days of housing t(36)=7.56, p=6.09e-09 and t=5.61(36),p=8.53e–7). Points are individual mice colored by genotype.

**Figure S7:**
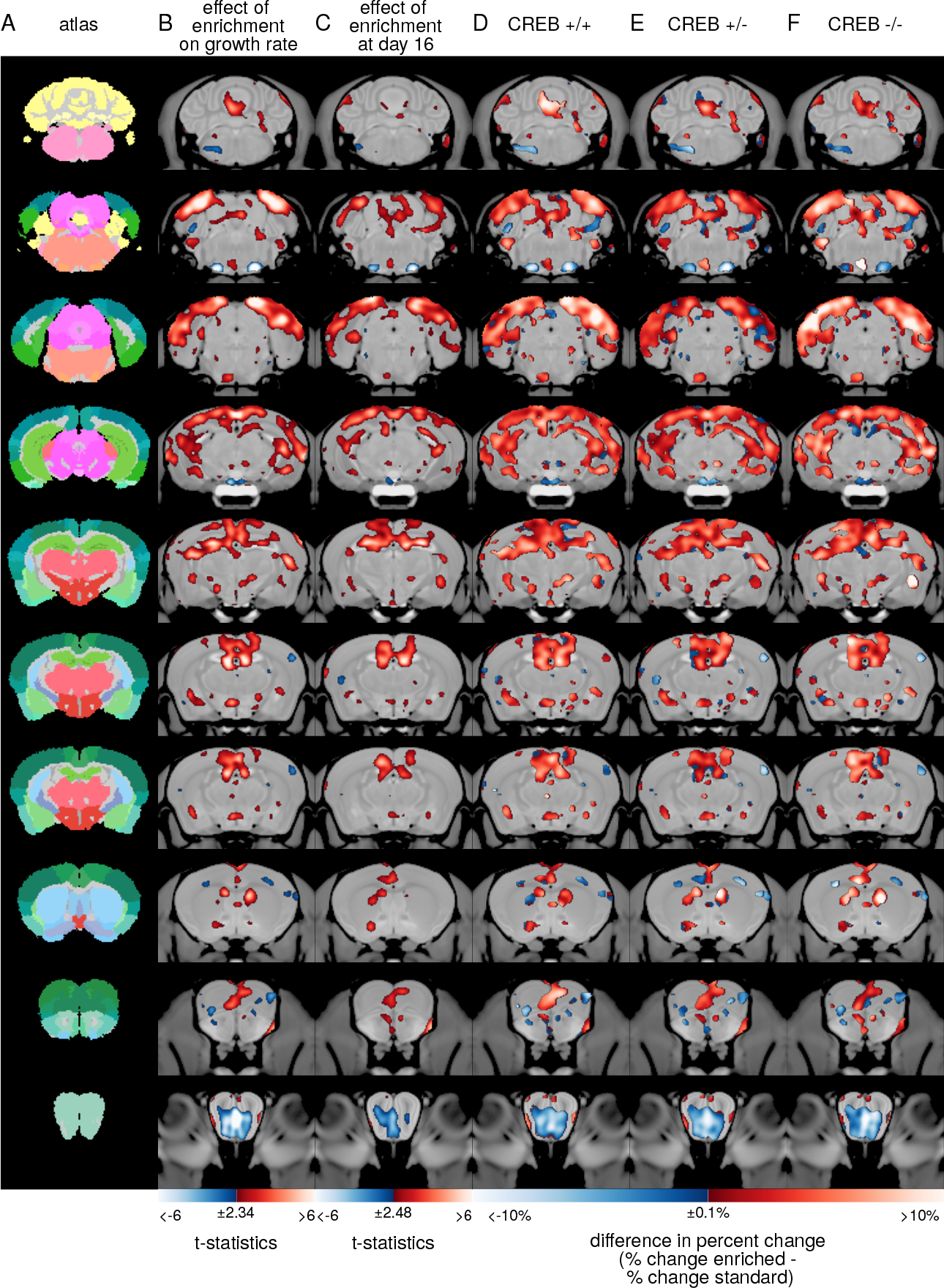
Voxel-wise analysis showing that loss of CREB does not affect volume changes induced by environmental enrichment. Coronal images taken from the average anatomical MR image of all mice in the study are shown overlaid with A) atlas labels which delineate 159 regions in the brain; B) t-statistics showing voxels where there was a significant effect of enrichment on the linear component of growth rate (interaction between enrichment and day) for the model including all genotypes of mice; C) t-statistics showing a significant effect of enrichment at day 16 of housing; D-F) maps showing the difference in percent change between enriched and standard housed wildtype (CREB*αδ*+/+) mice (D), CREB*αδ*+/− mice (E), and CREB*αδ*−/− mice (F) at each voxel. Only voxels where there was a significant effect of housing condition at day 16 or a significant housing-day interaction at a 10% FDR are shown. Red colours indicate voxels where enriched mice grew at a faster rate or had a larger volume after 16 days when compared to standard housed mice. Blue colours indicate voxels that grew at a slower rate or had smaller volume after 16 days relative to standard housed mice. Sample sizes are shown in table S1.

**Figure S8:**
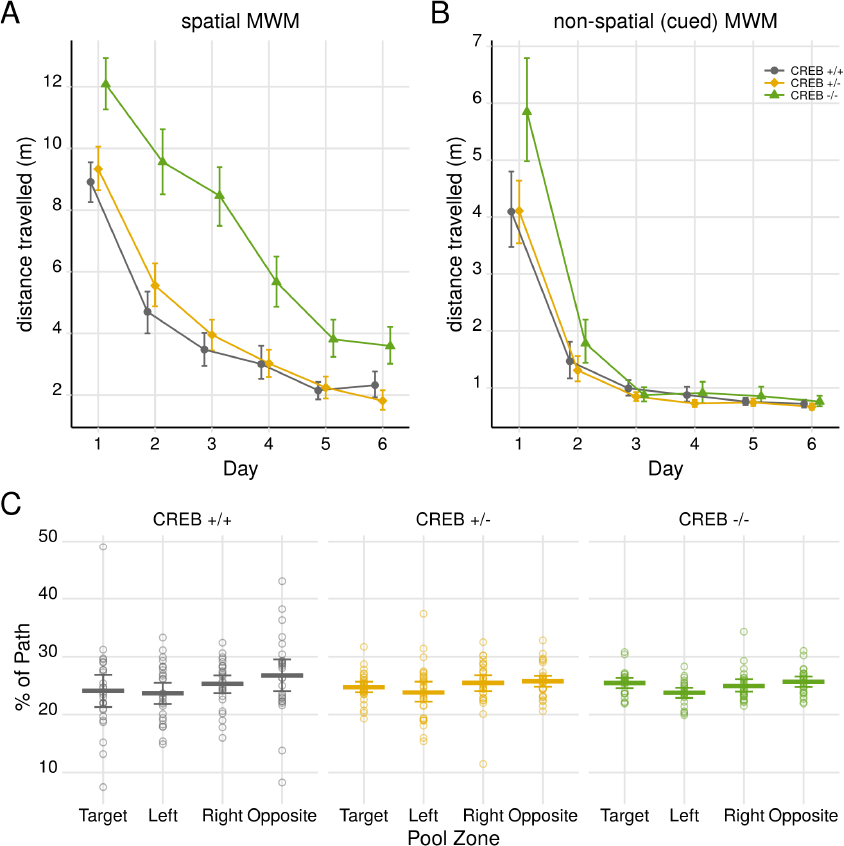
Loss of CREB impairs spatial memory. A) CREB*αδ*−/− mice have impaired learning on the spatial version of the Morris Water Maze and learn at a slower rate than CREB*αδ*+/− and CREB*αδ*+/+ mice (interaction between day and genotype: χ^2^(8)=65.268,p=4.27e–11, interaction between day and CREB*αδ*−/−: t=−5.53,p=3.5e–8). After 6 days of training CREB*αδ*−/− mice travelled significantly further than wild types to locate the platform (t(168)=3.327, p=0.001). B) CREB*αδ*−/− mice have intact non-spatial learning. CREB genotype only subtly altered the rate at which mice learned the non-spatial version of the MWM (interaction between day and genotype: χ^2^(8)=53.23 p=9.74e–09). CREB*αδ*−/− mice travelled further to the platform on day 1 (t=4.805,p=2.19e–06) but showed no differences after day 1. C) Non-spatially trained mice show no preference for platform location. Points represent group means. Error bars are 95% confidence intervals.

**Figure S9:**
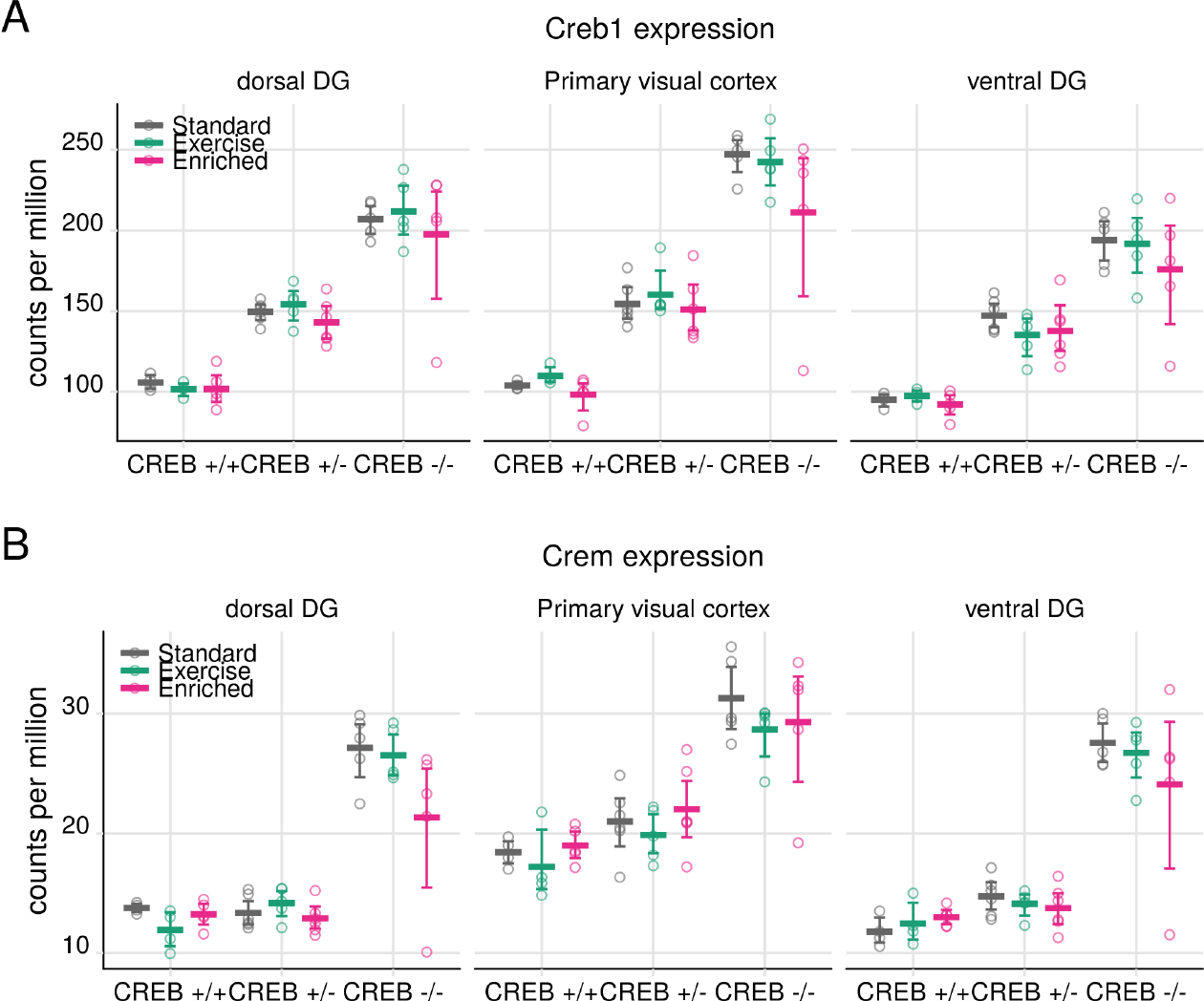
CREB mutants display compensatory upregulation of CREB and CREM. Loss of CREB*αδ* is associated with compensatory upregulation of CREB*β* and CREM, which leads to significant upregulation of total Creb1 (A) and Crem (B) in CREB mutant mice in all brain regions sampled. Points are individual mice. Lines are group means. Error bars are 95% confidence intervals. DG = dentate gyrus.

**Figure S10:**
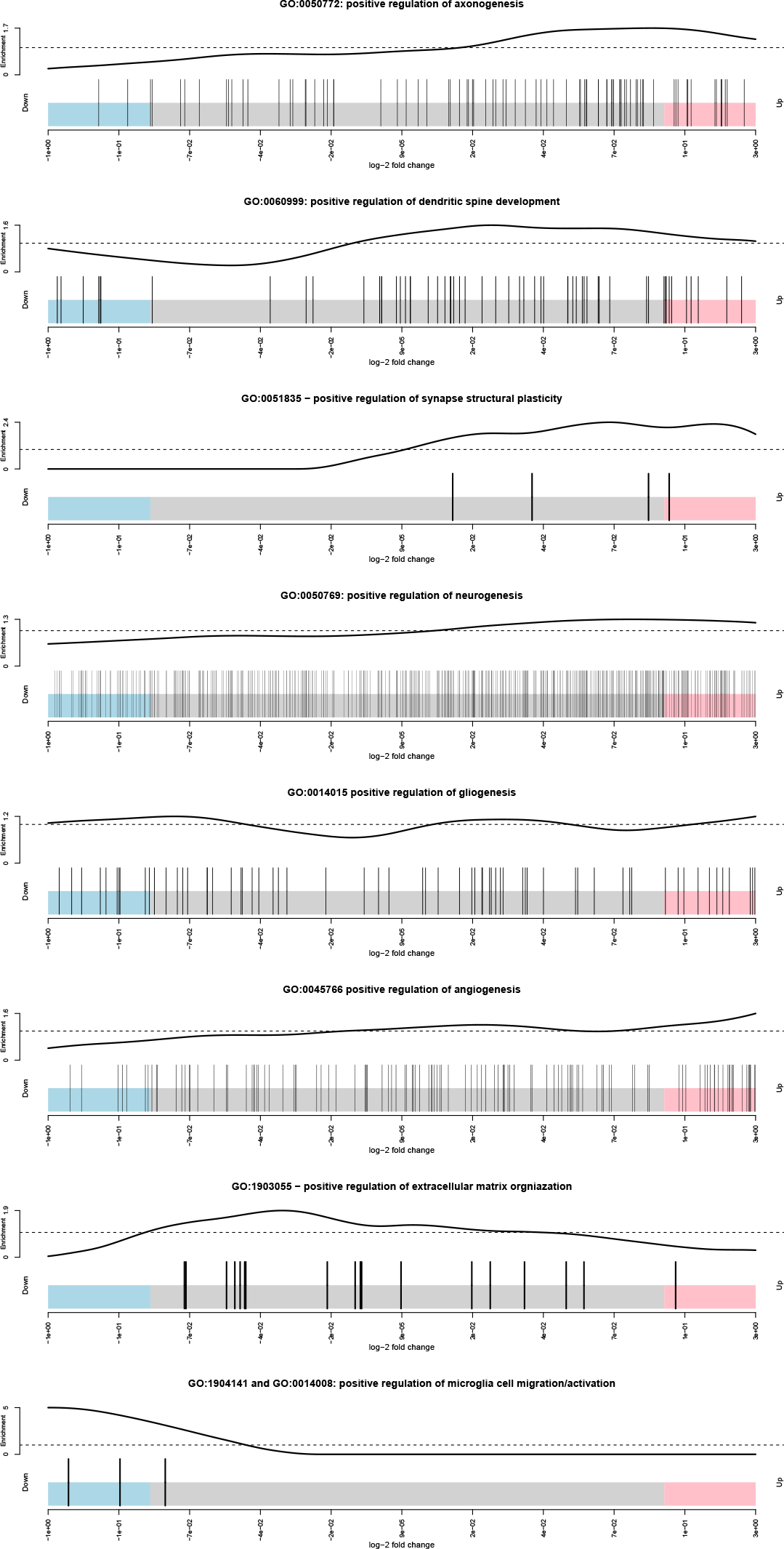
Environmental enrichment is associated with upregulation of genes associated with positive regulation of neurogenesis and neuronal re-modelling and with differential expression of genes involved with angiogenesis. Each barcode plot shows the log-2 fold change of genes associated with a given gene ontology (GO) term. Genes are shown as vertical bars which look similar to a barcode and are ranked from left to right by increasing fold change. A log-2 fold change of 9e–05 corresponds to a fold change of 1 in enriched versus standard housed mice (i.e. no change). The top and bottom 10% of differentially expressed genes are shown in pink and blue, respectively. The curve above the barcode shows the relative enrichment of genes in each part of the plot. For example, genes associated with GO:0060999 tend to be upregulated while those associated with GO:1904141 or GO:0014008 (microglia migration/activation) tend to be downregulated.

**Figure S11:**
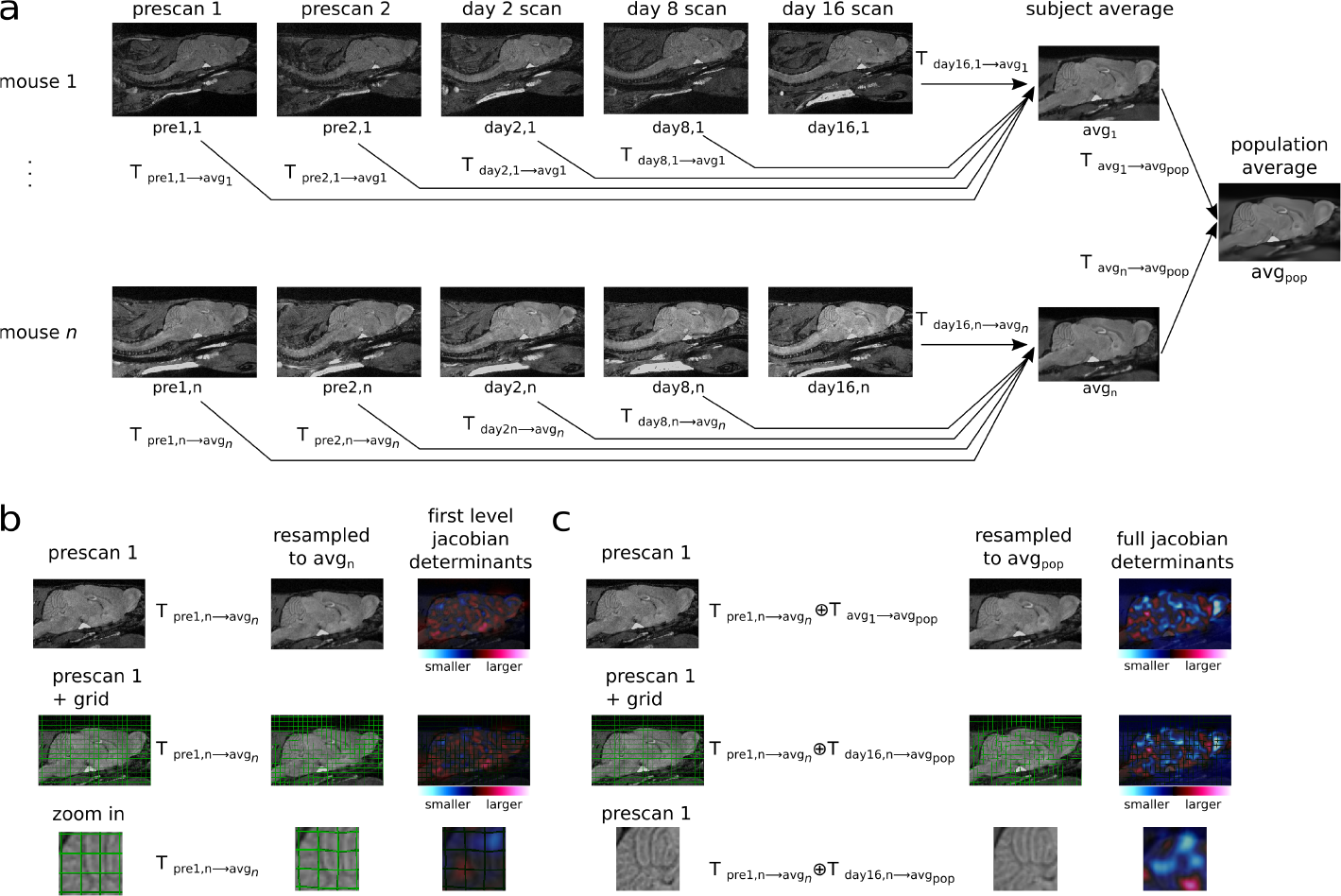
Illustration of image registration process. a) The image registration is performed in two stages. First, for each mouse, the images from each timepoint are registered together to generate a subject-specific average. Each input image is related to the subject average by a transformation *T*. Then the subject averages are aligned to generate a population average (*avg_pop_*). The input images are related to the population average by the concatenation of two transforms: the transformation between the input image and the subject average and the transform between the subject average and the population average. Concatenation is shown by ⊕. b) Applying the transformation shown to the input image (top) resamples the input into the same space as the subject average. The deformations can be visualized in the middle and bottom rows. Input images are shown overlaid with gridlines which become warped upon transformation to the subject average (middle). Bottom row shows voxels in the cerebellum deforming towards the average. Volumetric changes can then be calculated by computing the Jacobian determinant of the inverse of this transformation (encoded by displacement fields). The Jacobian determinant represents volume expansion or shrinkage at each voxel. For example, a Jacobian determinant value greater than 1 would indicate the voxel is larger than the subject average while values less than 1 indicates the voxel is smaller than the subject average. The Jacobian determinants calculated from the transformations relating the input images to the subject specific averages are called “first level Jacobians” and were used to analyse growth rates. c) The input images can also be aligned to the population average by concatenating the transforms. The Jacobian determinants calculated from this concatenated transformation are called “full Jacobian determinants”.

**Figure S12:**
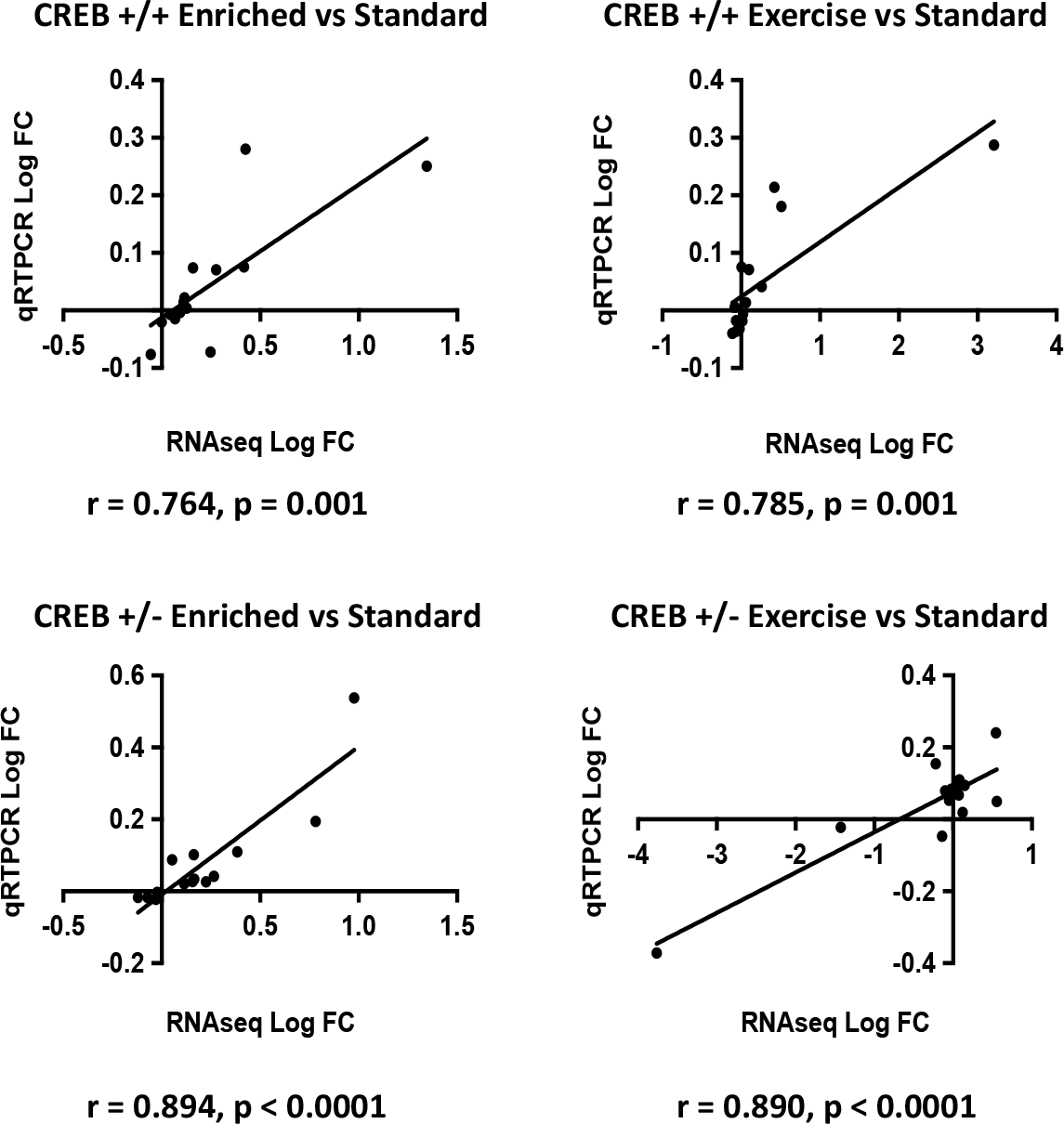
qRTPCR validation of RNAseq data. RNAseq data was validated using qRTPCR of 15 selected genes across the three brain regions analyzed (dorsal dentate gyrus, ventral dentate gyrus, and visual cortex). Plots show correlation between log fold change (FC) in expression of each gene, as computed with RNAseq and qRTPCR data. There was a significant correlation between RNAseq and qRTPCR data in all comparisons.

### 10.3 Supplementary Tables

**Table S1:** Number of mice used for enrichment and exercise experiments.

| Sex | Genotype | Standard Housing | Isolated Standard Housing | Exercise | Enrichment |
| --- | --- | --- | --- | --- | --- |
| Female | CREB$\alpha\delta$+/+ | 12 | 9 | 8 | 14 |
| | CREB$\alpha\delta$+/- | 10 | 7 | 8 | 11 |
| | CREB$\alpha\delta$-/- | 8 | 7 | 10 | 9 |
| Male | CREB$\alpha\delta$+/+ | 19 | 8 | 11 | 16 |
| | CREB$\alpha\delta$+/- | 24 | 5 | 10 | 24 |
| | CREB$\alpha\delta$-/- | 18 | 7 | 12 | 17 |

**Table S2:** Number of mice used for water maze experiments.

| Sex | Genotype | Control | Non-Spatial | Spatial |
| --- | --- | --- | --- | --- |
| Female | CREB$\alpha\delta$ +/+ | 16 | 12 | 10 |
| | CREB$\alpha\delta$ +/- | 13 | 14 | 15 |
| | CREB$\alpha\delta$ -/- | 16 | 13 | 12 |
| Male | CREB$\alpha\delta$ +/+ | 14 | 14 | 14 |
| | CREB$\alpha\delta$ +/- | 14 | 16 | 14 |
| | CREB$\alpha\delta$ -/- | 12 | 11 | 13 |

**Table S3:** Number of mice used for RNA sequencing

| Region | Genotype | Standard | Exercise | Enrichment |
| --- | --- | --- | --- | --- |
| dorsal DG | CREB$\alpha\delta$ +/+ | 4 | 4 | 5 |
| | CREB$\alpha\delta$ +/- | 6 | 5 | 6 |
| | CREB$\alpha\delta$ -/- | 5 | 5 | 5 |
| ventral DG | CREB$\alpha\delta$ +/+ | 4 | 4 | 5 |
| | CREB$\alpha\delta$ +/- | 6 | 5 | 6 |
| | CREB$\alpha\delta$ -/- | 5 | 5 | 5 |
| visual cortex | CREB$\alpha\delta$ +/+ | 4 | 4 | 5 |
| | CREB$\alpha\delta$ +/- | 6 | 5 | 6 |
| | CREB$\alpha\delta$ -/- | 5 | 5 | 5 |

**Table S4:**
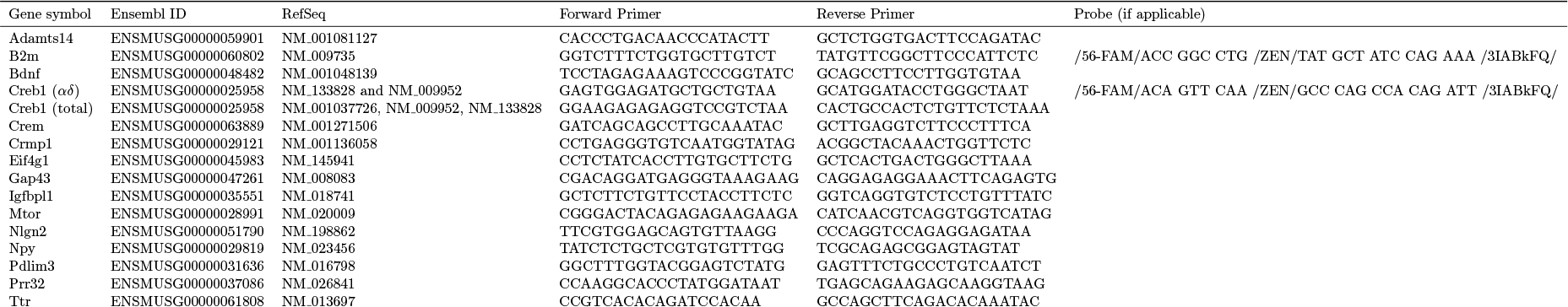
Genes and primers used for qRTPCR validation of RNAseq data.

**Table S5:**
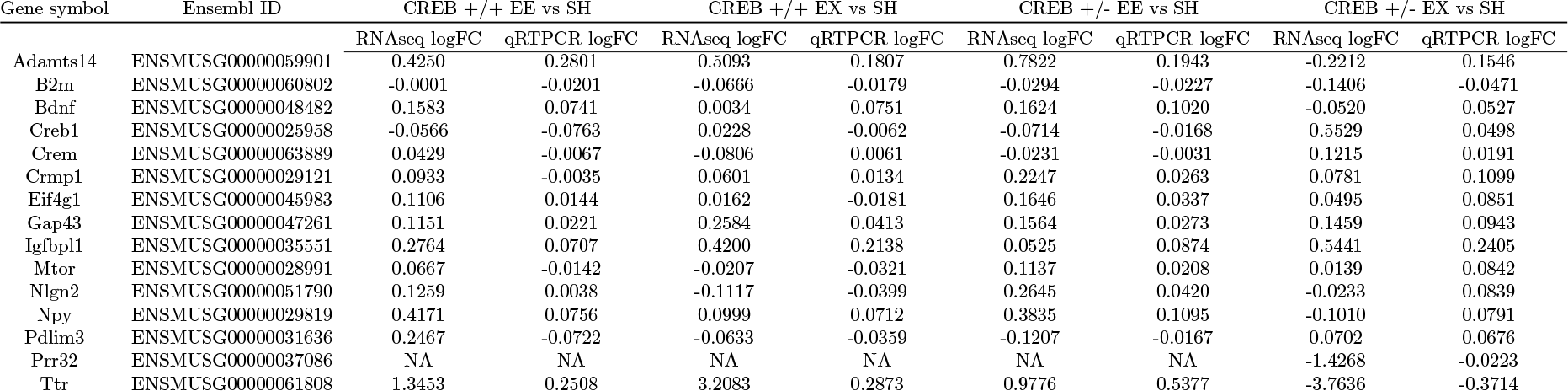
Log fold change for 15 genes of interest as computed using transcriptional data acquired with RNAseq and qRTPCR. NA values are reported for genes in which cycle threshold value in the qRTPCR assay was greater than 30, indicating that expression is low and unreliable for small sample sizes.

